# Structural basis for activation and mechanism of the bacterial hexameric motors RadA and ComM in DNA branch migration during homologous recombination

**DOI:** 10.1101/2024.02.15.580443

**Authors:** Leonardo Talachia Rosa, Emeline Vernhes, Anne-Lise Soulet, Patrice Polard, Rémi Fronzes

## Abstract

Some DNA helicases play key roles in genome maintenance and plasticity through their branch migration activity in distinct pathways of homologous recombination. RadA is a highly conserved bacterial helicase involved in both DNA repair in all species and in natural transformation in Gram positive firmicutes. In others, ComM is the natural transformation-specific helicase. Both RadA and ComM form hexameric rings and use ATP hydrolysis as a source of energy. In this work we present the cryoEM structures of RadA and ComM interacting with DNA and ATP analogues. This is the first structure reported for ComM, and the first structure of RadA in its active state. In particular, this study unveiled a molecular switch for ATP hydrolysis and DNA binding coupling in RadA and the role of the Lon protease-like domain, shared by RadA and ComM. Together our results provide new insights on the branch migration mechanism at the molecular level in the context of homologous recombination.

## Introduction

Homologous recombination (HR) is a conserved mechanism of DNA strand exchange that is central to genome biology in all living cellular organisms (bacteria, archaea and eukaryotes) and in some of their viruses. Many different HR pathways are key to common and essential cellular processes such as DNA repair and replication, meiosis in eukaryotes, or horizontal gene transfer in prokaryotes. Any failure in these pathways can have dramatic consequences, threatening the viability or genetic integrity and evolvability of the cell, which can lead to many pathological disorders such as cancer or developmental defects (Kreuzer, 2005; Sun et al., 2020).

HR pathways proceed through successive steps, with some reactions common to all and others more specific. The initial HR steps consist of DNA strand exchange reactions that are invariably catalyzed by a conserved ATPase known as the HR recombinase, called RecA in bacteria, RadA in archaea, and Rad51 and Dmc1 in eukaryotes. This enzyme acts in the form of a polymer initially assembled on single-stranded DNA (ssDNA), known as the presynaptic HR filament. This first HR intermediate is capable of searching for a complementary sequence in a recipient double-stranded DNA (dsDNA) molecule. The recombinase then proceeds to pair the bound ssDNA molecule with its complementary sequence in the dsDNA, creating a branched DNA molecule (Hertzog et al., 2023; Yang et al., 2020). This second HR intermediate is called the synaptic or heteroduplex HR product and is most commonly referred to as the D-loop structure. Each HR pathway involves a distinct subset of effectors that act with or independently of the recombinase to orchestrate the formation of the presynaptic and synaptic HR intermediates and the D-loop maturation steps. A D-loop processing reaction common to many HR pathways is the subsequent pairing of the second DNA strand of the donor and recipient DNA molecules, on one or both sides of the D-loop, leading to the formation of a single or double Holliday junction (HJ or dHJ, respectively). The final outcome of each HR pathway is then determined by how these three- or four-stranded HR intermediates are processed. A key reaction is to extend or reverse the exchange of paired DNA strands by driving DNA branch migration, either at one or both boundaries of the D-loop or at each HJ. DNA synthesis may be involved in the maturation step of these different HR products. Finally, another known and distinct maturation step of these HR intermediates consists in their specific cleavage to resolve and separate the joined DNA molecules (Sun et al., 2020; Yang et al., 2020).

A distinct D-loop maturation reaction that drives three-stranded DNA branch migration has recently been characterized in bacteria in the HR pathway of natural transformation (NT). This reaction is catalyzed by either RadA (not to be confused with archaeal recombinase) or ComM ATPases, depending on the species. NT consists of the uptake and internalization of exogenous DNA into the cell as linear ssDNA, which is then integrated into the genome by RecA-directed HR. This key biological process alters the genetic content of bacteria in a variety of ways. These include gene insertion and deletion, as well as multiple types of genomic rearrangements, all of which promote bacterial adaptation and evolution in response to stress, including antibiotic resistance and vaccine escape (Johnston et al., 2014). In contrast to RadA, which is widely conserved across bacterial species, ComM is less well distributed, as notably absent in Gram positive firmicutes (Nero et al., 2018). RadA is universally involved in HR-dependent DNA repair and is additionally involved in NT in the model transformable firmicutes species *Streptococcus pneumoniae* and *Bacillus subtilis* lacking ComM (Beam et al., 2002; Torres et al., 2019). In contrast, ComM has been shown to be exclusively involved in NT in other transformable species, including the gram negative species *Vibrio cholerae* (Nero et al., 2018). Comprehensive biochemical characterization of RadA from *Escherichia coli*, *S. pneumoniae*, *B. subtilis* and *Thermus thermophilus* and ComM from *V. cholerae* revealed that they are functional homologs during HR (Cooper & Lovett, 2016; Marie et al., 2017; Torres et al., 2019; Inoue et al., 2017; Nero et al., 2018). Both RadA and ComM were found to be ring-shaped hexameric DNA motors that direct three-stranded DNA branch migration in vitro and are powered by ATP hydrolysis. In addition, genetic analysis of their central role in NT of *S. pneumoniae*, *B. subtilis* and *V. cholerae* clearly demonstrated that both promote elongation of ssDNA integration in this HR pathway.

RadA and ComM differ significantly at the structural level. RadA is composed of an N-terminal C4 zinc finger domain (ZF domain), a central RecA-like ATPase domain, and a C-terminal Lon protease-like domain (Lon domain) (Figure 1a). RadA from *S. pneumoniae* forms hexamers in vitro in the absence of DNA or ATP and translocates on DNA in the 5’ to 3’ direction, as deduced from its helicase activity on various DNA substrates (Marie et al., 2017). In parallel with its biochemical analysis, we reported the crystal structure of *S. pneumoniae* RadA in complex with dTDP (Marie et al., 2017). In this structure, RadA forms a hexameric ring with C2 symmetry. While the Lon and ATPase domains were resolved, the ZF domain was not visible. The ATPase active sites were all in an inactive conformation. The atomic structure of ComM is not known, but it is predicted from sequence analysis to contain an N-terminal Lon protease-like domain (Lon domain), a central AAA+ ATPase domain with an embedded zinc finger, and a C-terminal Mg^2+^ chelatase domain (Figure 1a). ComM was described as a monomer in solution, forming hexamers in vitro only in the presence of ATP and DNA, and was shown to have bidirectional helicase activity (Nero et al., 2018), unlike RadA from *S. pneumoniae*.

**Figure 1:**
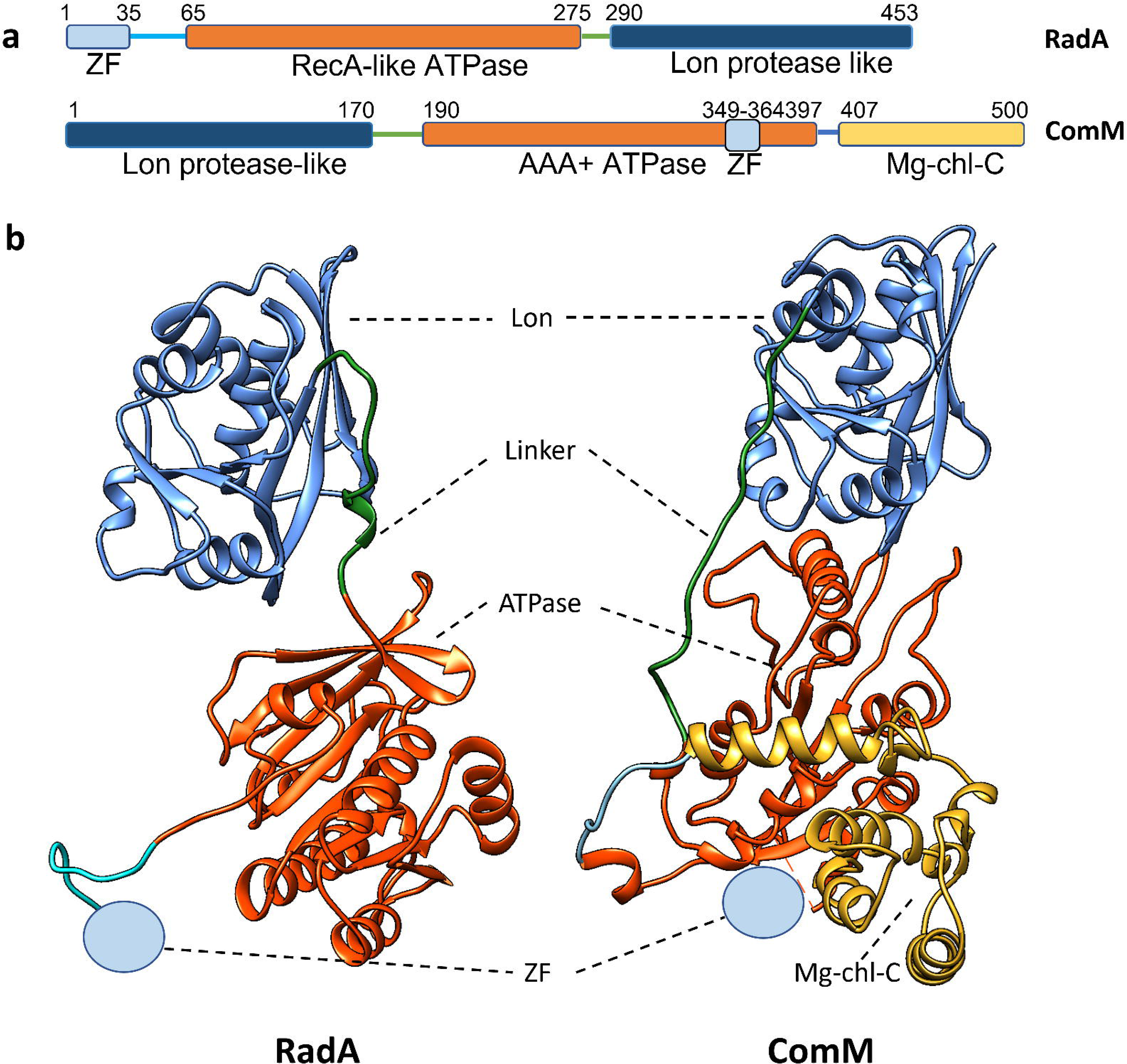
Domain representation of RadA and ComM. (a) Sequence representation of RadA (top) and ComM (bottom), separated by domains with respective starting and ending residues of each domain labeled on top. Lon protease-like domains are represented in dark blue and ATPase domains (RecA-like and AAA+) in orange. Linkers connecting the Lon and ATPase domains are in green. Predicted C4 metal binding domains are in light blue. C-terminal Magnesium-chelatase-like domain is depicted in gold. (b) Structure of RadA (left) and ComM (right) monomers, represented as ribbons and coloured accordingly to the sequence representation in (a). Subunit C of the active hexamers obtained by cryoEM was chosen for both proteins, represented with the same Lon domain orientation. The C4 metal-binding domain is not resolved in either structure and is depicted as a light-blue circle.

While the role of RadA and ComM in DNA branch migration during HR is well established, little is known about the molecular basis of their interaction with DNA, including their functional assembly and ATP-dependent translocation on DNA. Here, we present and compare the cryoEM (cryogenic electron microscopy) structures of RadA and ComM from *S. pneumoniae* and *Legionella pneumophila*, respectively. Both proteins were found to be stably bound to dsDNA in the presence of non-hydrolyzable ATP analogues, revealing the overall architecture of these two translocases in a catalytically active state. While RadA and ComM share the same hexameric scaffold formed by the Lon domains, they show substantial differences in the arrangement of the ATPase domains relative to the DNA and the Lon domain ring. These findings shed light on different mechanisms of ATPase activation by DNA and the regulatory function of the Lon domain in these two hexameric motors that drive three-stranded DNA branch migration during HR.

## Results

### Cryo-electron microscopy reveals the active conformation of RadA bound to DNA

RadA from *S. pneumoniae*, purified from *E. coli* mainly as a hexamer, was shown to exhibit ATPase activity in the presence of DNA in a previous study (Marie et al., 2017), with lower activity on dsDNA than on ssDNA. To gain a deeper understanding of the interaction between RadA and DNA, purified RadA was incubated in the presence of ATP-y-S, a poorly hydrolyzable ATP analog, with a gapped DNA substrate consisting of 30 base pairs of double-stranded DNA followed by 60 nucleotides of single-stranded DNA, and an additional 30 base pairs of double-stranded DNA (Combination CD in Supplementary Table 1). Density maps of hexameric RadA bound to DNA were obtained using cryoEM, yielding a resolution of 3.15 Å (Figure 2a– left panel and Supplementary Figure 1). The central panel of Figure 2a presents the atomic model, demonstrating an overall protomer structure that closely resembles the RadA crystal structure (PDB: 5LKM), which we previously solved in the absence of DNA (Marie et al., 2017). In the crystal structure, the central ATPase domains (residues 65-275) and the C-terminal Lon domains (residues 290-453) of each subunit of the hexamer are connected by a 15-residue linker (residues 275-290). The N-terminal ZF domain could not be resolved.

**Figure 2:**
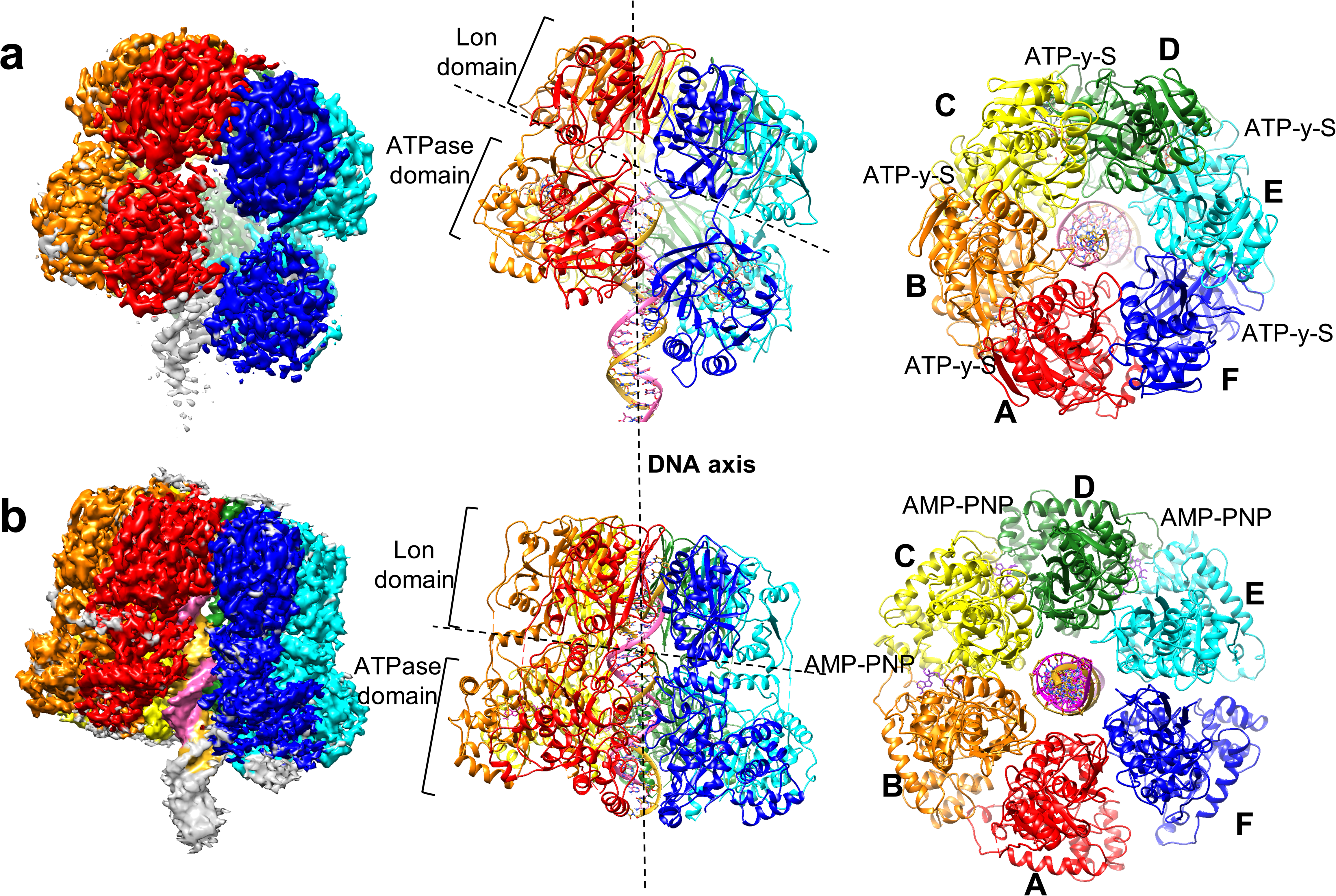
CryoEM analysis of DNA-bound helicases RadA and ComM. (a) CryoEM analysis of DNA-bound RadA. The final 3.15 Å map at level 0.01 (left panel) is colored according to the molecular model (central panel). RadA is arranged as a hexameric ring, with the Lon domains in a planar conformation, and the ATPase domains orchestrated in a helical spiral, accompanying the 21-bp double-stranded DNA in the middle, which goes from the limit between Lon and ATPase domains in direction of the N-terminal region, until density fades out. RadA is tilted in relation to the DNA axis, where the Lon axis is estimated to be in a tilt of 25°. The RadA hexamer is colored by chain in rainbow representation, from the most retracted ATPase domain (Chain A-red) to the most stretched (Chain F-blue) in relation to the Lon domains. Leading DNA strand is colored in magenta, and lagging strand in golden. On the view from the C-terminal Lon domain (right panel), ATP-y-S labels are marked in between adjacent ATPase domains where the density map indicates presence of the ligand. (b) CryoEM analysis of DNA-bound ComM. The presented map (left panel) is a composite map, at level 0.1, of 3 different local refinement maps (Lon domains at 2.8 Å, trimer composed by monomers C-E+DNA at 3.23 Å, and whole hexamer, at 3 Å). The combined map is coloured according to the molecular model (central panel). ComM is arranged as a hexameric ring, with both the Lon and ATPase domains in a planar conformation, orchestrating the 22-bp double-stranded DNA in the middle, which goes from the limit of the Lon ring N-terminal aperture and crosses the central pore, until density fades out in the C-terminal region. ComM has a small tilt in relation to DNA, estimated at 7°. The ComM hexamer is coloured by chain in rainbow representation following the RadA colouring (Chain A-red to Chain F-Blue). Leading DNA strand is colored in magenta, and lagging strand in golden. AMP-PNP labels are positioned in the view from the N-terminal Lon domain (right panel) where the density map indicates presence of ligand.

In the RadA cryoEM structure, the arrangement of the Lon domain remains consistent with the crystal structure of RadA without DNA. The Lon domain ring exhibits C6 symmetry, while the ATPase domains are arranged in a helical configuration in the hexamer (Figure 2a - middle panel). The flexible linker between the Lon and ATPase domains allows for this asymmetry mismatch (Figure 1b). Five of the six ATPase domains (Subunits A-E) are clearly defined in the density map. Subunit F, which is located farthest from the Lon domain, exhibits greater flexibility in certain areas (Supplementary Figure 1). As in all RecA-like ATPases, the ATP binding site is located between adjacent domains. The map shows that five molecules of ATPγS and one Mg^2+^ ion are well positioned within the active site (Figure 3a). In the cavity between subunits A and F, only a fuzzy density was visible, indicating partial nucleotide binding in this region. Each protomer comprises an N-terminal arm that encircles the adjacent ATPase domain on subunits A-E, as previously observed in the crystal structure (Figure 2a- right panel). The N-terminal ZF domain was not visible in the map (Supplementary Figure 1), suggesting that it is highly flexible relative to the ATPase domain, even when bound to DNA.

**Figure 3:**
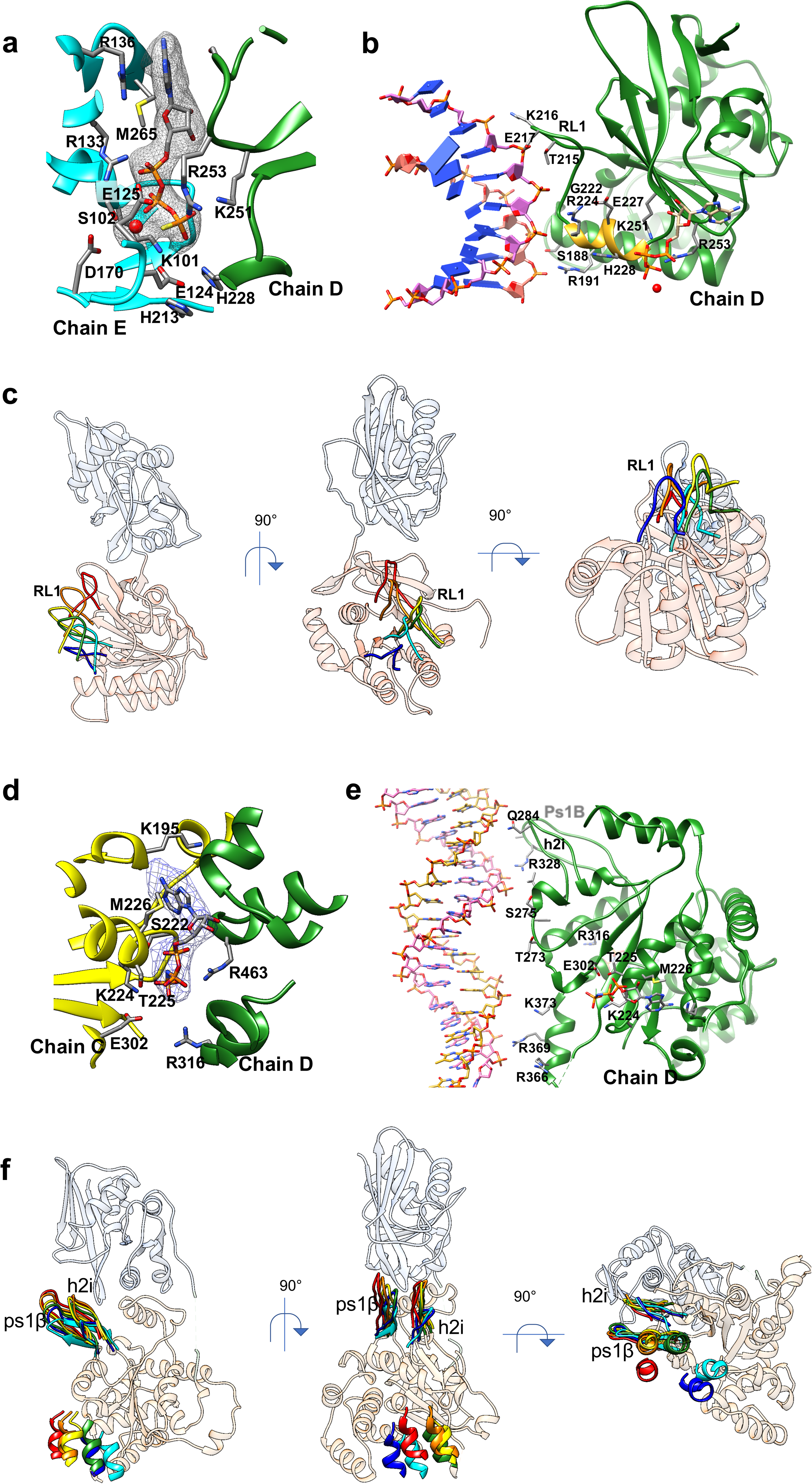
Cooperativity between DNA and ATP coordination in RadA and ComM. (a) ATPase pocket of RadA bound to ATP-y-S, between protomers D and E. ATP-y-S is shown in stick representation colored by element, inserted in its density depicted as a grey mesh. Mg^2+^ ion is depicted as a red sphere. Protein is in ribbon representation colored by chain: Chain D – Green; Chain E - Cyan. Relevant residues are depicted as sticks and colored by element. (b) DNA Binding in RadA. ATP coordination leads to the structuration of the short helix (in gold) on the N-terminal of the KNRFG motif, pushing the DNA binding loop forward to interact with the DNA leading strand (magenta). Cleavage of ATP results in loop retraction. (c) Movement of RadA DNA binding loops along the DNA translocation process. Superposition of DNA binding loops for the six subunits in RadA, with monomer C shown as transparent ribbons, with the Lon Domain in blue and the ATPase domain in orange. Only the DNA binding loops are shown for the other subunits, following the color-code in figure 2. Different angles allow visualization of movement in the Y axis (left panel), X axis (central panel) and Z axis (right panel). (d) ATPase pocket of ComM bound to AMP-PNP, between subunits C and D. AMP-PNP is shown in stick representation colored by element, inserted in its density depicted as a blue mesh. Protein is in ribbon representation colored by chain: Chain D – Green; Chain E - Cyan. Relevant residues are depicted as sticks and colored by element. (e) DNA binding in ComM, orchestrated by two loops: pre-sensor 1 beta-hairpin (Ps1B-residues 324-336) and Helix-2-insert (h2i-residues 281-289). Additional contacts occur via a helix at the C-terminus of the C4 domain (residues 366-373) (f) Movement of ComM DNA binding loops along the DNA translocation process. Alignments and coloring obey the same parameters used for RadA (c).

A clear density of a 21-base-pair dsDNA is visible. The density begins within the RadA hexamer at the boundary between the Lon and ATPase domains and expands towards the ATPase domain and beyond the hexamer ring (Figure 2a - left panel). As the DNA becomes less restricted and more flexible, the density of the DNA progressively decreases. RadA appears to bind to the dsDNA ends of the gapped DNA substrate with its ssDNA extending to the outside of the hexameric ring on the ATPase side. While the DNA backbone has a similar structure to that of DNA-B (supplementary figure 2), the base pairing in the downstream region (towards the Lon domain) seems to be distorted or disrupted.

### CryoEM structures of *Legionella pneumophila* ComM

We overexpressed ComM from *L. pneumophila* in *E. coli*, attaching a C-terminal His6-tag. The protein was purified through nickel affinity chromatography and eluted as a single peak on a Superdex-200 10/300 gel filtration column. We observed a 50 kDa monomer protein state (see Supplementary Figure 3). This is in agreement with the results reported for the homologous ComM protein from *V. cholerae*, which was shown to hexamerize only in the presence of ATP and ssDNA (Nero et al., 2018). Here, we incubated purified *L. legionella* ComM protein with AMP-PNP and a synthetic oligonucleotide consisting of 60 bp of dsDNA and 60 nt of ssDNA (Supplementary Table 1 - Combination EG), which enabled us to obtain the cryoEM density map of ComM hexamers bound to DNA and AMP-PNP at a resolution of 3.1 Å (Figure 2b and Supplementary Figure 3). Local refinements were also carried out on half of the hexamer and the Lon domain, resulting in an improvement in the quality of the map in these regions (Supplementary Figure 4a-c).

The final model of the ComM protomer (Figure 1b) reveals that the N-terminal domain does comprise a Lon protease-like domain, as deduced from its sequence. Its AAA+ ATPase domain is similar to those of archaeal and eukaryotic MCM/CMG helicases and bacterial proteins such as RavA and magnesium chelatase (Supplementary Figure 5) (Baretić et al., 2020; Fodje et al., 2001; Jessop et al., 2020). Additionally, the C-terminal Mg^2+^ chelatase domain is closely linked to the ATPase domain (Figure 1b). Densities for the zinc finger domain of ComM were visible in the cryoEM map (Figure 2b), but the resolution was too low for accurate modeling. Notably, although ComM and RadA have an inverted sequence order, they share identical topology between the ATPase and Lon domains (Figure 1b). The long linker (residues 170-190) between the domains in ComM enables domain swapping. Without its ZF domain, the ComM hexamer consists of a double ring (Figure 2b). One ring is formed by the Lon domains and the other by the ATPase domains, which are surrounded by Mg^2+^ chelatase domains (which do not interact with each other in the ring). The ATPase ring remains nearly planar relative to the Lon domain, except for the DNA binding loops (as presented below in the ‘ DNA binding to ComM’ section). The density that corresponds to a 24 bp long DNA segment was distinctly visible throughout the hexamer. This DNA follows the hexamer symmetry axis, in contrast to the tilted axis seen in RadA (compare Figure 2a and Figure 2b).

Data processing of ComM revealed the dataset could be divided into three populations. 53% of the particles correspond to the helicase bound to DNA and AMP-PNP, used to generate the structure described above, and presumably stabilized in a translocated state. In 23% of the dataset, ComM hexamers are observed in the absence of DNA (Supplementary Figure 6). The ATPase domains adopt a cracked-ring C2 symmetry in this structure, while the Lon domain remains unchanged in comparison to the active conformation. This energetic state may represent either a transient DNA pre-loading or post-release complex.

In the remaining 24% of the particles, ComM hexamers interact *via* their ATPase domains to form a dodecamer that coordinates a single molecule of dsDNA (Supplementary Figure 7). Previous studies on MCM members of the Superfamily 6 of helicases have repeatedly reported the formation of dodecamers as pre-activation complexes (Cheng et al., 2022; Miller et al., 2019; Noguchi et al., 2017).

### ATPase active site in RadA

In our previous study (Marie et al., 2017), we attempted to obtain an active state of RadA by crystallizing it in the presence of dTTP. However, the nucleotide was hydrolyzed during the crystallization process, and as a result, the structure was obtained in the presence of dTDP. Here, we reveal a cryoEM structure of the RadA hexamer bound to dsDNA with five ATPase active sites occupied by ATP-y-S and the last unoccupied (Figure 2a and Figure 3a). Using the D/E subunit interface as a model, R136 and M265 in subunit E and G255 and S256 in subunit D coordinate the positioning of the adenosine base in ATP-y-S. This occurs at the end of the KNRFG motif (residues 251 to 255), a conserved motif unique to RadA homologs (Marie et al., 2017). The Walker A motif (residues 95 to 103) in subunit E coordinates the alpha and beta phosphates with R133. Sterically constrained on the opposite side is R253 of the KNRFG motif in subunit D, serving as an arginine finger to promote energy conservation during catalysis (Figure 3a). Other several interactions stabilize the gamma-phosphate, including K251 from the KNRFG motif in subunit D. This specific residue is commonly termed as the "piston" for ATP cleavage and its mutation significantly hinders transformation *in vivo* (Marie et al., 2017). The catalytic glutamate, E125, in subunit E coordinates a nucleophilic water molecule for ATP hydrolysis (Lyubimov et al., 2011). Further coordination is achieved by G100, K101, and S102 from the Walker A motif of subunit E. This motif additionally coordinates with a Mg2+ ion, as well as with E124 and D170 of subunit E (Figure 3a). This description is in agreement with our previous results (Marie et al., 2017), which showed that mutants in the Walker A (K101A) and KNRFG (K251A and R253A) motifs were severely affected for ATPase activity.

### ATPase active site in ComM

The ATPase domains of the bacterial chaperone protein RavA (PDB 6SZB) and the archaeal MCM replication helicase (PDB 6MII) are very similar to the ComM ATPase domain (Supplementary Figure 5). The structure of ComM that is bound to dsDNA was obtained in the presence of the non-hydrolyzable ATP analog AMP-PNP, which is located at the interface between two adjacent subunits (Figure 3d). Densities for three AMP-PNP molecules were observed at the interfaces between chains B and E, involving four protomers (see Figure 2b right panel).

Taking the interface between chains C and D as a model, the adenosine nucleotide is coordinated by M226 and K195 of subunit C (Figure 3d). The beta-phosphate is surrounded by residues 221-225 of subunit C Walker A (residues 218 to 225). The gamma-phosphate is coordinated by the side chains of K224 (lysine piston), T225, and E302 of subunit C. Additionally, the side chains of R316 (sensor 3 motif) and R463 (arginine finger) of subunit D are also involved in the coordination.

### DNA binding to RadA

In the crystal structure of RadA without DNA, the ATPase domains organize in a planar hexameric ring (Marie et al., 2017). Upon interaction with dsDNA, the ATPase domains adopt a helical organization. This places the KNRFG motif, which includes both the arginine finger and the catalytic piston, in close proximity to the bound ATP molecule of the neighboring subunit. At the bottom of the ATP-binding pocket, the terminal phosphate is bound by H213 in subunit E and H228 in subunit D (Figure 3a). This location is consistent with the positions of H465 and Q494 in the T7 Gp4 helicase, a close structural homolog of RadA, which coordinates the binding of a nucleophilic water molecule with ATP hydrolysis to facilitate DNA binding (Supplementary Figure 2) (Gao et al., 2019). In RadA, the stacking of these two histidine structures a short helix at the N-terminus of the KNRFG motif (R224 to H228), which in turn pushes the adjacent RL1 loop (RadA loop 1, residues 213 to 223) forward to promote DNA binding (Figure 3b). RL1 includes DNA binding regions found in the D1 and D2 loops of the T7-gp4 structure but lacks the extended loop region found in its homologue (Supplementary Figure 2d), a distinctive feature that is likely compensated for by additional DNA contacts with the Lon domain region (Inoue et al., 2017). Indeed, three residues (R305, R314, and K345) in the Lon domain of RadA from *T. thermophilus* (RadAT.t) (corresponding to K320, N329, and K360 in RadA from *S. pneumoniae*, as shown in Figure 4f) are located on the inner surface of the Lon ring and may directly mediate DNA binding (Inoue et al., 2017).

**Figure 4:**
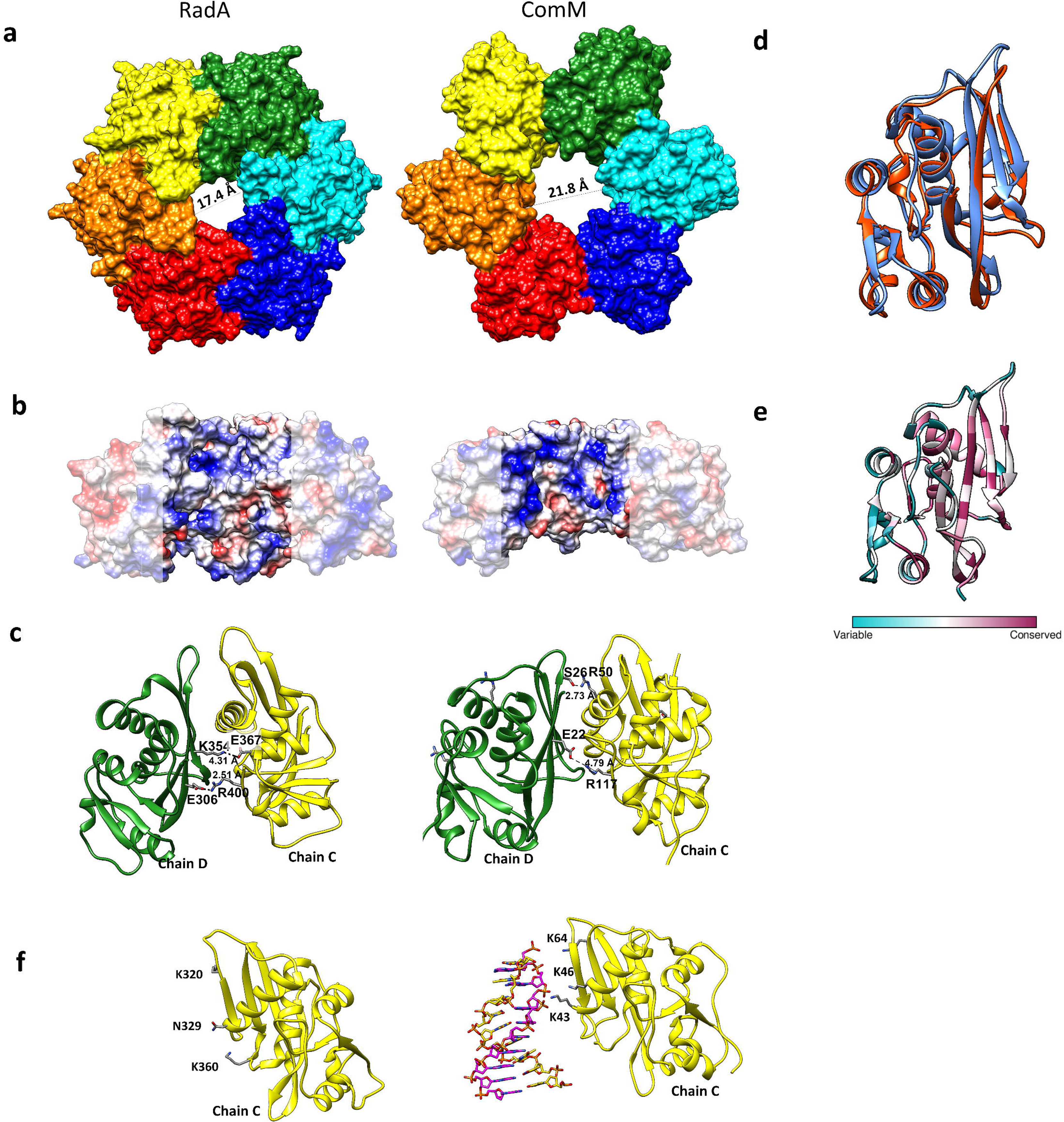
Comparison of the Lon domains of RadA and ComM. (a) Lon domain surfaces for RadA (left) and ComM (right), colored by chain. RadA structure shows an aperture on the external end of 17.4 Å, while ComM shows a larger aperture of 21.8 Å. (b) Columbic potential of a trimer of Lon domains, allowing for visualization of the center of the channel. The Lon half-ring is shown at 90° in relation to (a). The sides of the subunits are white-shaded, while the interior of the ring remains framed in the middle. Columbic potential is depicted by color, from negative (red) to neutral (white) and positive (blue). (c) Interface between Lon subunits C and D on RadA (left) and ComM (right), highlighting the main salt bridge and hydrogen bonds participating in his interface, with respective distance between them (d) Superposition of Lon domain monomers of RadA (orange-residues 291 to 452) and ComM (blue – residues 5 to 168), showing an RMSD of 0.987 Å over 87 pruned atoms. (e) Consurf analysis of RadA Lon domain, with a color gradient depicting variable (cyan) to conserved (magenta) residues. (f) Charged residues pointing towards the center of the pore in the Lon domain of RadA (left) and ComM (right), potentially involved in the coordination of DNA. For ComM, the DNA is also shown in Magenta and golden, while for RadA no defined density is observed in this region.

The primary contact with DNA is established by K216, which binds to the phosphate of the DNA backbone and coordinates the leading 5’ to 3’ DNA strand in a manner similar to that of DnaB loops (ZF to Lon domain polarity)(Gao et al., 2019). Additionally, T215, E217, and G222 are involved in coordination (see Figure 3b). A second loop named RL2, which consists of residues 170 to 189, coordinates the lagging 3’ to 5’ strand. S188 and R191 attach to the DNA backbone six base pairs below the base pair coordinated by RL1 towards the ZF domain, when the DNA helix completes a half turn on its axis and the other strand is exposed to the corresponding RadA subunit (see Figure 3b and Supplementary Video 1).

The ATPase domains’ helical ring topology mirrors the DNA topology. RL1 and RL2 interact with two nucleotides above their adjacent domain (from Lon to ZF). Subunit A binds with DNA downstream (towards the Lon domain), while subunit E binds with DNA upstream. (see Figure 3c and Supplementary Video 1). RL1 in subunit F is slightly displaced and seems dissociated from DNA, confirming that ATP hydrolysis results in DNA release.

### DNA binding to ComM

Although the ATPase domains of ComM also assemble in a subtle helical pattern, they remain almost planar in relation to the Lon domain. In MCM members of the helicases superfamily 6, DNA binding occurs through two conserved hairpins directed towards the center of the pore. These are the pre-sensor I beta hairpin (Ps1B) and the helix 2 insert (H2i) (see Figure 3e and Supplementary Figure 5). The flexibility of these hairpins enables them to arrange themselves helically on DNA, creating an uninterrupted hydrophobic chamber while leaving the rest of the domain undisturbed (Fletcher et al., 2003). ComM maintains the Ps1B loop from residues 324 to 336, with R328 forming a crucial hydrogen bond with the DNA phosphate backbone and the side chains of A329 and A330 creating a hydrophobic surface (see Figure 3e and Supplementary Figure 5). H2i, which consists of residues 281 to 289, is half the size of its MCM counterpart and prevents the extended hydrophobic interaction along the DNA, instead making contact *via* a hydrogen bond with Q284 (Figure 3e). Both loops follow the helical pattern of the DNA and bind 2 nucleotides apart from the previous subunit, resulting in the binding of 12 nt per cycle (Figure 3f and Supplementary Video 2). In the coordination of the lagging strand, S275 makes hydrogen bond contacts with the phosphate backbone, while the remaining residues in loop 273-277 form a hydrophobic surface. A second region of the protein via R366, R369, and K373 residues makes further contact with the lagging strand. (See Figure 3e, Supplementary Video 4, and Supplementary Figure 8 for more information.)

### RadA and ComM share the same Lon-protease-like scaffold

RadA and ComM both contain a Lon protease-like domain, although they belong to different ATPase families. In RadA, the domain is situated in the C-terminal region, whereas in ComM it is located in the N-terminal region (Figure 1a). Due to the extended linker that connects the Lon and ATPase domains in ComM, the topology of the Lon in relation to the ATPase domain is conserved in both proteins. Despite their low sequence identity (28%), which is mostly located in the ß-sheet core (Figure 4e), the two Lon domains superimpose with a root mean square deviation (RMSD) of only 1 Å (Figure 4d). In both DNA-bound proteins, as well as in the RadA crystal structure (PDB 5LKM) and in DNA-free ComM (Supplementary Figure 6), the Lon domain forms a rigid hexamer (Figure 4a) with similar pore diameters (17.4 Å in RadA and 21.8 Å in ComM). The use of the PDB-PISA server reveals a significant difference in the buried surface area between adjacent subunits of the two proteins, with ComM measuring 678 Å^2^ and RadA at 1028 Å^2^ (Supplementary Table 2). The interface between subunits displays polar contacts that may aid in stabilizing the hexamer arrangement. RadA shows electrostatic interactions between E306 and R400, as well as K354 and E367. In ComM, there are electrostatic interactions between S26 and R50 as well as E22 and R117, as shown in Figure 4c.

In contrast to traditional Lon proteases, the interior of the Lon domain ring is rich in basic residues, consistent with potential DNA coordination at the center of the pore (Figure 4b). This is further supported by the observation that ComM interacts with DNA via three lysines, K43, K46 and K64, which form hydrogen bonds with the DNA backbone within the central pore (Figure 4f). Consistent with this, it has been reported that the Lon domain of RadA from *T. thermophilus* displays DNA-binding activity and that mutations in three residues directly facing the center of the pore, K320, N329 and K360 in RadA from *S. pneumoniae*, alter the DNA-binding ability of full-length RadA (Figure 4f)(Inoue et al., 2017).

### In RadA, a molecular switch between the Lon and ATPase domains is critical for coupling ATP hydrolysis to DNA binding

In *T. thermophilus* RadA, two surface residues, R286 and R385 (R301 and R400 in *S*. *pneumoniae* RadA), play a critical role in the DNA-mediated ATPase and DNA-binding activities (Inoue et al., 2017). R400 is located at the interface between the RadA Lon domains, as discussed above. The role of R301 has been unclear. This residue is positioned at the interface between the Lon and ATPase domains. It was hypothesized that this residue could regulate the ATPase domain of RadA, rather than directly affect DNA binding.

While the ATPase domains adopt a helical conformation in the DNA-bound RadA hexamer, the Lon domains remain planar. Thus, R301 interacts with different regions of the ATPase domains depending on the topology of each subunit (Figure 5a). In subunits A and B, where the ATPase domain is located closer to the Lon domain, R301 appears to form a salt bridge with E239. The density map suggests an alternative occupancy in which the arginine would not make a contact between the domains. Meanwhile, in subunits C to E, where the ATPase domains are farther from the Lon domain, the density map clearly shows an interaction between R301 and E76 (Figure 5a). However, in subunit F, which is unbound to DNA, the ATPase domain is too distant to contact R301. Thus, we propose that R301 functions as a salt bridge switch, differentiating and helping to stabilize the different conformational states of the ATPase domain.

**Figure 5:**
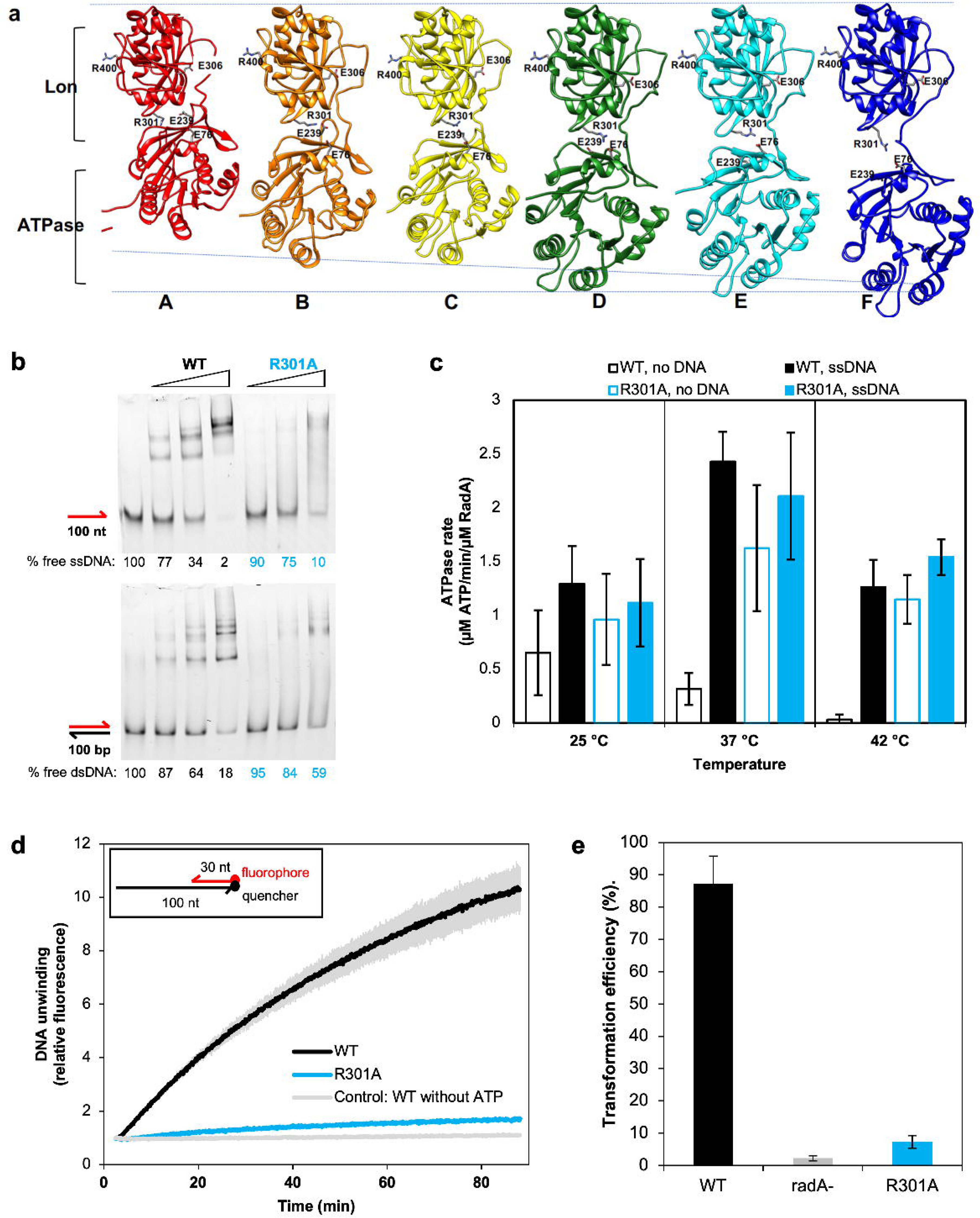
Salt-bridge switch during RadA activity. (a) RadA protomers are represented side-by-side in equivalent orientation. Protomers color follow the same scheme as in figure 2. Along the translocation cycle, the ATPase domain moves farther from the Lon domain, accompanied by a subtle rotation movement. This rotation results in a salt bridge switch formed by R301, in the Lon domain, and E239 (monomers A,B and C) and E76 (monomers D and E), in the ATPase domain. (b) DNA-binding properties of RadA mutants by Eletrophoretic Mobility Shift Assay (EMSA). Binding to single-stranded DNA is shown in the upper panel, while binding to double-stranded DNA is shown in the lower panel. Free DNA migration position is indicated by the respective DNA depiction on the left (fluorescent DNA in red, unlabeled DNA in black), while protein-bound DNA will have a delayed migration. DNA is incubated with incrementing quantities of protein, according to the gradient bar shown on top. Quantification of the free DNA bands is shown under each gel as a percentage of the first lane (no RadA) band. (c) ATPase activities at 25, 37 and 42 °C, in the absence or presence of single-stranded DNA. Error bars are calculated from at least three experimental replicates. (d) Helicase activities. DNA unwinding is shown as fluorescence increases over the initial fluorescence. A scheme of the DNA substrate used for the helicase assays is presented in the inset. The curves represent the mean of two experimental replicates and error bars are shown as faded colors. (e) *In vivo* transformation efficiencies in *S. pneumoniae*. Error bars are calculated from at least five experimental replicates.

To evaluate the functional role of this molecular switch, we attempted to purify site-directed alanine mutants of residues R301 and E239, which form this bridge in RadA from *S. pneumoniae*. The R301A site-directed mutant of RadA was successfully purified as wild-type RadA in the form of a soluble hexamer, but the E239A mutant was found to precipitate immediately upon purification, precluding its biochemical analysis. Therefore, we studied only the R301A mutant, starting with the analysis of its DNA binding properties by electrophoretic mobility shift assay (EMSA), as previously done with wild-type RadA (Figure 5b)(Marie et al., 2017).

Figure 5b shows that RadA_R301A_ has a significantly lower affinity for both ss- and ds-DNA. This suggests that the RadA binding to DNA is dependent on the stabilization of the ATPase domain through the R301/E76-E239 bridge. The ATPase activity of the R301A mutant was then analyzed in the absence and presence of ss-DNA. The R301A mutant exhibited high levels of basal ATPase activity in the absence of DNA, indicating that ATP hydrolysis is decoupled from its interaction with DNA and ATP is consumed in a futile cycle (Figure 5c). This effect was partially mitigated when the experiments were conducted at a lower temperature (25°C instead of 37°C), but it was still evident at higher temperatures (42°C). The previous sections describe the thermal effect on the coupling between ATP hydrolysis and DNA binding. To investigate the decoupling between ATP cleavage and DNA binding of the RadA_R301A_ mutant, we analyzed its helicase activity. The assay involved hybridizing a 30 nt long DNA strand labeled with a fluorophore at its 5’ end to the 3’ region of a 100 nt strand labeled with a quencher at its 3’ end. The spatial proximity of the two labels causes fluorescence quenching, resulting in low fluorescence intensity. When the two strands dissociate due to RadA helicase activity, the labels move apart, and fluorescence increases (Figure 5d). Consistent with the ATPase and DNA binding assays, the RadA_R301A_ mutant exhibits very low helicase activity (Figure 5d). Finally, we investigated the impact of disrupting the salt bridge between the Lon and ATPase domains of RadA on bacterial transformation activity (Figure 5e). The RadA_R301A_ mutant strain exhibited a 92% reduction in efficiency, comparable to the *radA*-null mutant (Marie et al., 2017; Peter et al., 2007).

Taken together, these results highlight an important role for the salt bridge switch mediated by R301 of the Lon domain and E76/E239 of the ATPase domain in the catalytic ATP hydrolysis cycle of RadA modulated by DNA and inferred to reflect its translocation on DNA powered by ATP hydrolysis. Disruption of the salt bridge can cause futile ATP consumption and alter DNA binding. This may be due to increased flexibility in the ATPase domain, preventing RadA from stabilizing the retracted or stretched conformations between the two domains. As a result, the RadA_R301A_ mutant can still coordinate ATP binding and hydrolysis, but cleavage occurs before DNA binding is promoted and the helical assembly is established. Breaking this bridge deactivates the protein’s DNA translocation and helicase activities, as well as its DNA branch migration activity on HR intermediates during NT.

## Discussion

In this study, we solved the structure of the DNA-bound ring-like hexameric helicases RadA and ComM, highlighting key features of their translocation mechanism on DNA. This structural analysis revealed that these two HR bacterial effectors have a modular and similar topology, organized as two-tiered hexamers, with one tier consisting of the ATPase domain ring and the second tier consisting of a Lon-like domain ring. Despite the inverted primary sequences of these domains, *i.e.* the Lon domain is located at the C-terminus in RadA and at the N-terminus in ComM, both share a similar topology thanks to a long linker that spans the entire length of ComM and spatially connects the two domains.

### The Lon domain acts as a common hexamerization module in RadA and ComM

Both RadA and ComM contain a Lon-like structural domain (Figure 4). This domain has not been found in any other DNA-binding ATPases characterized to date, suggesting that it may confer specific properties to these two helicases in the context of HR.

For RadA, it was previously proposed that the Lon domain acts as a scaffold for its hexamerization, which occurs in the absence of DNA and ATP (Marie et al., 2017). On the other hand, ComM hexamerization was mainly observed in the presence of ATP and DNA, and to a lesser extent in the presence of ATP alone, as reported for ComM from *V. cholerae* and confirmed here for ComM of *L. legionella* (Nero et al., 2018). The PDB PISA server displays a significantly different interaction surface between the Lon domains in ComM and RadA. RadA has a greater number of residues involved in interfacial interaction, resulting in a larger interfacial area and greater solvation free energy upon hexamerization compared to ComM (−6.8 kCal/mol for RadA vs. −3.9 kCal/mol for ComM) (see Supplementary Table 2). Additionally, the p-value linked to the free energy gain for RadA of 0.28 indicates a site-specific interaction between subunits. Conversely, the p-value for ComM, which is close to 0.5, reflects a non-specific interaction and is similar to the average value for its structure. These findings suggest that the hexamerization of RadA, which is independent of DNA and ATP, is primarily driven by global solvation free energy rather than by interactions between specific residues.

### The Lon domain functions as a clamp on DNA

Ring cracking is a widely accepted mechanism for loading hexameric helicases onto DNA (Fernandez & Berger, 2021). The eukaryotic replicative MCM1-6 helicase (SF6) is known to self-load onto DNA through a gap between the MCM2 and MCM5 subunits of the hexamer. It then self-assembles at the origin of replication as a pair of hexamers linked by their N-terminal regions. Phosphorylation by additional factors culminates in activation and dodecamer disruption (Miller et al., 2019). The hexameric SF4 DnaB replicative helicase of *E. coli* requires the DnaC protein to break the ring and mediate its DNA loading (Arias-Palomo et al., 2019). RadA and ComM are both capable of self-loading on DNA, with RadA mostly in the form of a pre-assembled hexamer and ComM in the form of a DNA-free monomer. Recent cryoEM studies by Shin et al. (2020) have shown that Lon proteases form a stable hexamer in the presence of protein substrate, but form a ‘lock washer-like’ cracked ring in the absence of substrate to facilitate substrate recognition and loading. Based on the structural similarity between the Lon protease domain and the Lon-like domain of RadA, it is proposed that the Lon domain promotes ring cracking in the presence of DNA to facilitate hexamer loading.

### RadA and ComM translocate on DNA in a helical arrangement

Both RadA and ComM hexameric structures encircle the DNA with poorly hydrolyzable ATP analogues, corresponding to their active translocation state on DNA. They exhibit a similar hand-to-hand DNA-binding mechanism. In RadA, the entire ATPase domain is arranged in a helical pattern around the DNA (Figure 2a). The DNA structure facilitates proper positioning of the ATP-binding pocket between two monomers (Figure 3a). When ATP binds, the KNRFG motif, which includes the Arginine Finger and Lysine piston that coordinate ATP hydrolysis, is repositioned. This, in turn, pushes the RL1 loop towards the DNA for interaction (Figure 3b). As demonstrated in the case of the T7-gp4 helicase (Gao et al., 2019), the hexamer’s ATPase domains exhibit a helical pattern, followed by a slight domain rotation that results in ATP cleavage in the most extended subunit relative to the Lon domain (Figure 3c).

We propose that, upon ATP hydrolysis, the KNRFG motif, which coordinates the γ-phosphate of ATP, would become non-structured, affecting the adjacent residues and retracting the loop from DNA binding. The ATPase domain is then free from the constraints caused by ATP and DNA, and retracts towards the Lon domain. Here, it coordinates a new ATP molecule and engages in a position 12bp upstream of the previous location. This culminates in a translocation cycle of 12 bp/6 ATP (see Supplementary video 3). In a previous study, we demonstrated that disrupting the Walker A motif by inserting a K101A mutation in RadA did not significantly impact DNA binding. However, introducing a K251A or R253A mutation, both in the KNRFG motif, resulted in reduced DNA binding (Marie et al., 2017). Based on the current model, this could be due to the inability of the KNRFG mutants to stabilize the necessary helical conformation required for DNA binding.

Building on previous site-directed mutagenesis work by Inoue et al. (2017), which found that mutations in different Lon domain residues affected DNA binding, we investigated the impact of R301 on RadA activity (R286 in RadA from *T. thermophilus*). In the CryoEM structure, R301 is involved in a salt-bridge switch with either E239 or E76 in the most retracted or stretched subunits, respectively (Figure 5 and Supplementary video 6). Our biochemical assays indicate that the disruption of this salt bridge results in futile ATP cleavage, even in the absence of DNA, and almost completely abolishes helicase activity (Figure 5c and d). *In vivo*, the R301A mutant exhibits a deficiency in natural transformation equivalent to that of a *radA* null mutant (Figure 5e). The salt-bridge between the Lon and ATPase domains may regulate RadA translocation activity by stabilizing either the retracted or stretched subunit forms and blocking ATPase activity in the absence of DNA.

In ComM, the helical arrangement is limited to the DNA binding regions, while the surrounding regions of the ATPase domain remain almost planar with respect to the Lon domain (Figure 2b). This is achieved in MCM proteins by two long flexible loops, named Ps1B and H2i, which form a continuous hydrophobic chamber for the translocation of dsDNA on the hexamer (Supplementary Figure 5)(Fernandez & Berger, 2021; Fletcher et al., 2003). Both loops are present in ComM, with Ps1B being highly similar but with H2i being much shorter. The H2i region typically includes an aromatic residue, such as Y386 in archaeal MCM or W569 in human MCM2, which intercalates between DNA bases. This interaction was proposed to either disrupt DNA stacking during strand separation or provide torque for translocation movement (Fletcher et al., 2003; Rzechorzek et al., 2020). The ComM H2i hairpin lacks an equivalent residue, but this may be compensated for by additional interactions between DNA and the Lon domain, which is absent in MCM homologues. The Lon domain of ComM contacts DNA at least three positions (Figure 4f).

In contrast to RadA, disruption of the Walker A motif in ComM results in a loss of DNA binding and helicase activity (Nero et al., 2018). Based on the topological arrangement of the ATP binding site in relation to the DNA binding loops, it is hypothesized that R316, which belongs to the sensor 3 motif, will advance the Ps1B hairpin upon ATP binding. Similarly, E302, located in the walker B motif, may have a comparable effect on the H2i hairpin, and R381 may have a similar sensing function that affects the binding of the 366-373 region. This results in a cooperative binding between DNA and ATP, as observed in RadA (Figure 3d and e). Disruption of ATP cleavage could cause conformational changes in the three sensor residues, resulting in the release of DNA.

### Dodecamerization of ComM

In 24% of the cryoEM data, two ComM hexamers interact back-to-back through their ATPase domains, coordinating a single dsDNA molecule that encompasses both hexamers (see Supplementary Figure 7). Dodecamers of the Superfamily 6 of helicases are well-known in the literature (Cheng et al., 2022; Miller et al., 2019; Noguchi et al., 2017). Eukaryotic and archaeal MCM helicases, involved in genome replication, dodecamerize upon recognition of the AT-rich region of the OriT to form a pre-activation complex at the initiation step of DNA replication. Once activated, the two MCM hexamers run past each other to perform strand separation (Miller et al., 2019; Noguchi et al., 2017). However, it is unclear whether the ComM dodecamer could act similarly during HR. This is due to the fact that the DNA encompassing MCM hexamers usually presents a short bending at the interface between the two hexamers, which serves as a base for forking initiation. This bending is not observed in ComM hexamers, where the dsDNA strongly maintains a classical DNA-B conformation along its entire length. The MCM proteins interact through their N-terminal domains (NTD) and move past each other, with the NTD in the front, in the 3’ to 5’ direction. Additionally, the NTD of MCM is topologically equivalent to the Lon protease domain of ComM. However, it appears that ComM dodecamerizes through the opposite interface, leaving the Lon domain at both extremities (see Supplementary Figure 7). This means that ComM hexamers would not move past each other upon activation of their translocation, but rather away from each other. Third, Unlike MCM, ComM is not involved in DNA replication initiation, but in the post-synaptic step of HR to promote DNA branch migration on the D-loop intermediate (Nero et al., 2018). One interesting possibility is that ComM would be recruited inside the D-loop, on the DNA heteroduplex resulting from RecA-directed ssDNA invasion and pairing. Within the D-loop, a single ComM hexamer may access its boundaries to facilitate homologous ssDNA incorporation in both directions by directing three-stranded DNA branch migration from these two sites.

### RadA and ComM as potential dsDNA translocases

This structural analysis of the interaction of RadA and ComM with DNA stabilized by poorly or non-hydrolyzable ATP analogues provides insight into how they might promote DNA branch migration during HR. Previous biochemical analyses of RadA proteins from different organisms have reported common but also some important functional differences in their DNA binding and helicase activities with respect to ATP binding and hydrolysis (Cooper & Lovett, 2016; Inoue et al., 2017; Marie et al., 2017; Torres et al., 2019). They were found to be ATPases activated by either ssDNA or dsDNA. However, they differ in their ability to promote helicase activity *in vitro*, which was observed for *S. pneumoniae* and *B. subtilis* RadA, but not for *E. coli* RadA (and not reported for *T. thermophilus* RadA). However, *E. coli* RadA, like *S. pneumoniae* and *B. subtilis*, promotes 3-strand DNA branch migration *in vitro* (Cooper & Lovett, 2016). Remarkably, 4-strand DNA branch migration has also been tested and observed for *E. coli* RadA (Cooper & Lovett, 2016). Our structural and functional analysis of *S. pneumoniae* RadA highlighted its homology with DnaB helicases and suggested that it could promote 3-strand DNA branch migration by translocating onto ssDNA at the D-loop boundaries in the 5’ to 3’ direction, coupling dsDNA strand separation with ssDNA pairing (Marie et al., 2017). Applying such a reaction at each D-loop boundary would favor ssDNA recombination in both directions, as supported by the genetic analysis of the role of RadA in NT (Marie et al., 2017). However, DnaB helicases are known to translocate along dsDNA and promote DNA branch migration by translocating along duplex DNA (Kaplan DL & O’Donnel M, 2002; Singleton et al., 2007). Such dsDNA encircling is exactly what was found in this structural analysis of RadA, which appeared to be stabilized in a translocation state with canonical DNA-binding loops interacting with one DNA strand (Figure 3).

Interestingly, biochemical analysis of ComM from *V. cholerae* showed that it exhibited bidirectional helicase and DNA branch migration activities (Nero et al., 2018). Similarly, the cryoEM structure of ComM from *L. legionella*, which we have characterized here, revealed its interaction with dsDNA.

These findings support a model in which these two hexameric DNA motors act as dsDNA translocases to direct 3-strand or 4-strand DNA branch migration. In this model of mechanism, RadA would be functionally similar to RuvB and RecG (Fernandez & Berger, 2021). This would explain the ATPase activity of RadA, which can be activated by either ssDNA or dsDNA, as well as why *E. coli* RadA does not exhibit *bona fide* helicase activity while being able to branch migrate double-stranded branched DNA substrate. This model raises two important questions. First, how could the Lon ring of RadA accommodate dsDNA? The average diameter of its positively charged pore is about 18 Å and is closed by flexible loops (residues 313-319) that define a constriction of 14 Å in diameter. B-DNA has an average diameter of 18 Å. Therefore, in the conformation we observed, the inner channel of the RadA Lon domain should be opened to allow the passage of dsDNA, possibly by pushing away these flexible loops. In contrast, we show that ComM has a larger pore radius with a diameter of 24 Å. This diameter is similar to the inner channels found in dsDNA translocases such as FtsK (Jean et al., 2020). Interestingly, the densities corresponding to DNA in this region are quite weak in ComM, suggesting a higher degree of flexibility and/or conformational heterogeneity for the DNA. A second question raised by this model is how these two DNA motors are recruited to the HR intermediates to promote DNA branch migration by translocating on dsDNA. Some clues are provided by their common key role in the extension of ssDNA incorporation from the HR D-loop of NT. This intermediate could not evolve in any other branched DNA structure because one of the two exchanged DNA molecules is the invading linear ssDNA. Thus, to prolong its recombination by translocating on dsDNA, these DNA motors could act by translocating the D-loop inwards on the duplex DNA assembled by RecA. At the 3-way DNA junction, the protein would couple dsDNA unwinding and ssDNA pairing to prolong recombination. Thus, RecA bound to dsDNA could define a landing pad for recruiting RadA and ComM inside the D-loop, giving them access to DNA for their translocation. In support of this model, physical and/or synergistic effects have been reported between RecA and RadA from *E. coli*, *S. pneumoniae*, and *B. subtilis* (Cooper & Lovett, 2016; Marie et al., 2017; Torres et al., 2019).

In conclusion, our structural and functional analysis of DNA-bound RadA and ComM provides important insights into how they act as DNA branch migration effectors during HR. These findings, combined with the previous biochemical analysis of these two hexameric motors that are widely conserved in bacteria, pave the way for future studies to uncover how they are recruited, loaded, and controlled in the HR pathways in which they specifically act, and how they mediate DNA strand exchange during their translocation.

## Materials and Methods

### RadA purification for CryoEM analysis

Full *radA* gene was amplified from *S. pneumoniae* genome and cloned in pASK-IBA3 vector, which adds a N-terminal strep-tag to the final protein product. After verification through sequencing, pASK-IBA3-*radA* was transformed into *E. coli* BL21 (DE3) for expression. The transformed strain was grown in 2 x 2L LB media in 5L conical flasks at 37 °C and 200 rpm until OD_595_ of 0.8, induced with 200 μg/L of anhydrotetracycline and incubated for 20 hours at 16 °C and 200 rpm. Cells were pelleted by centrifugation at 15 000 x g for 20 min at 4 °C and resuspended in 50 ml of lysis buffer (Hepes 50 mM pH 8.0; NaCl 100 mM; MgCl_2_ 2 mM) plus DNAse (5 µg/ml), Lysozyme 1 mg/ml and 1 tablet of EDTA-free protease inhibitor (ROCHE). Cells were broken passing twice through an Emulsiflex C5 (ATA scientific) and debris were pelleted by centrifugation at 100 000 x g for 20 min. The supernatant was loaded on a 5 ml Strep-tag purification column, washed with assay buffer (Hepes 50 mm pH 8.0; NaCl 100 mM) and eluted with 30 ml of assay buffer supplemented with 0.5 mg/ml Desthiobiotin (Sigma). Fraction containing RadA were pulled together and injected in a 5ml HiTrap heparin Column (GE) for DNA dissociation and eluted in assay buffer with a NaCl gradient ranging from 0.1 to 2 M. Protein presence was confirmed by electrophoresis in a 15% SDS-PAGE gel, run at 200 V for 40 minutes.

### ComM purification for CryoEM analysis

Full *comM* was amplified from *Legionella pneumophila* paris genome and cloned in pCDF-Duet1 to generate a C-terminal His-tagged ComM as a product. After verification by sequencing, pCDF-Duet1-*comM* was transformed in BL21 (DE3) for expression. The transformed strain was grown in 2 x 2L LB media in 5L conical flasks at 37 °C and 200 rpm until OD_595_ of 0.8, induced with 1 mM IPTG and incubated for 20 hours at 16 °C and 200 rpm. Cells were recovered, lysated and centrifuged as for RadA. The supernatant was loaded on a 5 ml His-trap column (GE), washed with assay buffer supplemented by 20mM Imidazole and eluted with assay buffer supplemented with 500 mM imidazole. Fractions containing ComM were pulled together, concentrated to 0.5 ml using a Vivaspin column with 30 kDa MWCO and injected in a Superdex 200 gel filtration column, run with assay buffer. ComM eluted as a single peak at 13 ml, consistent with a molecular weight of 50 kDa. Protein presence was confirmed by electrophoresis as for RadA.

### CryoEM sample preparation

Synthetic oligos were ordered from Eurofins. Hybrid DNA was prepared by mixing semi-complementary oligo-nucleotides (Supplementary table 1) at 100 µM final concentration of each in assay buffer supplemented with 1 mM DTT, boiled for 5 minutes and letting it cool down at room temperature. DNA-binding reaction for RadA was started by mixing 0.35 mg/ml RadA, 10 µM of hybrid DNA (Supplementary table 1-combination CD) and 5 mM ATP-γ-S, followed by a 30 minutes incubation at 37°C. DNA-binding reaction for ComM was started by mixing 1 mg/ml ComM, 10 µM of hybrid DNA (Supplementary table 1-combination EG) and 5 mM AMPP-N-P, followed by a 10 minutes incubation at 37°C. Quantifoil Cu300 mesh R2/2 grids were glow discharged at 0.3 mbar vacuum and 2 mA current for 35 seconds. 4 µl of reaction was added to the grid and blotted for 2.5 seconds in a Vitroblot instrument (FEI), at 100% humidity and 4 °C, at blot force 0, before being plunge-frozen in liquid ethane and transferred/stored in liquid nitrogen.

### RadA CryoEM Data collection and processing

Sample screening and pilot data collection was performed in a 200 kV Talos Arctica equipped with a K2 summit camera at the Institut européen de chimie et Biologie (Pessac-FR) using SerialEM (version 3.8) software. Final RadA Data collection was performed in a 300 kV Titan Kryos cryo-microscope (EFI) at ESRF (Grenoble-FR) (experiment MX-2261) at 165k magnification, pixel size of 0.827 Å, total dose of 54 e^-^/Å^2^ and 4.5 seconds exposition, using EPU software. A total of 8983 movies were collected and processed using Relion 3.1 software (Scheres, 2012). 2000 particles were manually picked and submitted to 2D classification in order to generate initial templates, which were in turn used for template-based picking. 908 107 particles were extracted and submitted to 3 rounds of 2D classification, resulting in 263 835 selected particles. The generated 2D classes were used as templates for an iterative round of template-based picking, which resulted in the picking and extraction of 3 768 530 particles. After several rounds of 2D classification, 1 258 631 particles were selected for a consensus 3D refinement without symmetry imposition, which generated a 3.54 Å overall resolution map, but in which only the Lon protease domains were well defined. This consensus map was used as a template for 3D classification without image alignment with a 4 class separation in order to further clean bad particles. 1 208 572 particles from good classes were selected, and submitted to a new consensus refinement, followed by a new 3D classification without image alignment in a 10-classes separation. The best class, containing 303 651 particles was selected and refined, generating a 3.6 Å overall resolution map. After per-particle CTF correction and Bayesian polishing, a final refinement was performed, generating a 3.15 Å overall resolution map. A scheme of the data processing is presented in Supplementary Figure 9.

The final map was post-processed using Autosharpen tool from Phenix package (Liebschner et al., 2019) using the two half maps in the input. Separated domains extracted from RadA’s crystallographic structure (Marie et al., 2017) were manually fit in the map using Chimera software, followed by a rigid body fitting using the RealSpaceRefine tool from phenix package(Liebschner et al., 2019). Manual curation and building of missing regions, as well as DNA and ATP-γ-S fitting was performed in *Coot*, followed by another round of RealSpaceRefine. Validation was first performed in *Coot* (Emsley et al., 2010) and subsequently checked using ‘CryoEM comprehensive validation’ tool from Phenix package(Liebschner et al., 2019).

### ComM cryoEM Data collection and processing

Sample screening and pilot data collection were performed as for RadA. Final ComM data collection was performed in a 300 kV Titan Kryos cryo-microscope (EFI) equipped with a Quantum-K3 camera at EMBL (Heidelberg – Germany) at 130k magnification, pixel size of 0.645 Å, total dose of 53.56 e^-^/Å^2^ and 1.2 seconds exposition, using SerialEM Version 3.8. A total of 32 534 movies were collected and processed in parallel using Cryosparc 3.3.1 and Relion 4.0 softwares (Punjani et al., 2017; Scheres, 2012). In Cryosparc, Movies were processed using PatchMotionCor and Patch CTF estimation. A subset of 2000 micrographs was used for blob picking with particle diameter of 100 Å. After 2D classification, 32 000 particles were selected and used as an input for picking using Topaz. After removal of duplicates, 1 687 279 particles resulted from Topaz picking, which were subjected to 2D classification, where 852 356 particles were selected. Data processing was twofold. In the first approach, the selected particles were subjected to ab-initial model generation with 10 classes. All the resulting models and particles were used as an input for a 10-class heterogeneous refinement. The promising classes were pulled together and submitted to a new round of ab-initial model generation, with 6 classes, and again, all the models were submitted to heterogeneous refinement with 6 classes. Two of these classes were selected, and submitted to unmasked refinement, followed by masked local refinement on the DNA-bound hexamer. This generated 2 subpopulations of DNA-bound ComM maps (main map EMDB 19574 PDB 8RXD, and subpopulation 2, EMDB 19575). In the second, classical approach, selected particles were subjected to ab-initial reconstruction with three classes. The best resulting model was used for a homogeneous refinement, resulting in an unmasked map of 3.17 Å, with well-defined Lon domain regions, but showing high variability on the ATPase region. This map was used as an input for a Non-Uniform refinement with a mask on the ComM hexamer, resulting in a hexamer consensus map of 3 Å (EMDB 19581). This consensus map was used as an input for a focused refinement on the Lon region, resulting in a 2.8 Å map for this region (EMDB 19578 PDB 8RXS). The hexamer consensus map was subjected to hexamer-masked hetero-refinement with 4 classes. The selected class contained 305 846 particles, and was subjected to a local refinement using a mask on half of the hexamer (a trimer) + DNA, resulting in a local map of 3.23 Å with considerably more homogeneity in the ATPase region (EMDB 19577 PDB 8RXK). A scheme of the data processing is shown in Supplementary Figure 9.

Initial molecular model for the ComM monomer was generated using Alphafold2 software (Jumper et al., 2021). The resulting model was divided into two domains (residues 1-180 and 181-500), copied 6 times to form a hexamer and manually fit in the main map using UFSC chimera (Pettersen et al., 2004). A 24mer dsDNA was generated in Chimera and fitted likewise. This model was submitted to initial adjustments in ISOLDE (Croll, 2018), followed by refinement in PHENIX RealSpaceRefine. Further adjustments were performed in *Coot* and ISOLDE before a final round of refinement and validation in PHENIX.

For the dodecamers, the 852356 particles from the consensus map were submitted to 3D classification with two classes, one of which showed initial density for ComM dodecamerisation, with 464 656 particles, but the second dodecamers had poor density due to high heterogeneity. After an unmasked 3D refinement, the subset was subjected to hetero-refinement with 10 classes. The best class, containing 69 029 particles, was refined, resulting in a 4.1 Å map of ComM dodecamers (EMDB 19580 - Supplementary Figure 9). Two ComM hexamer models were fit in the dodecamer density using UFSC Chimera, excluding the DNA chains. A 48mer dsDNA was generated in Chimera and fitted in the density, and further adjusted using ISOLDE.

For the DNA-free ComM hexamer, in Relion 4.0, Movies were processed using MotionCor2 (Zheng et al., 2017), with 10×10 patches, and Gctf (Zhang, 2016). Laplacian picking was used on a 1000 micrographs subset. After classification, 10 000 particles were used as input for Topaz picking on the whole dataset (Bepler et al., 2019), resulting in 2 888 884 picked particles. After rounds of 2D classification, 621 282 particles were selected and submitted to initial model generation. The initial model was refined without mask, resulting in a 6 Å consensus model. 3D classification with 10 classes and no particle alignment was performed, which revealed a class containing 78 222 particles of DNA-free ComM hexamers. This class was subjected to masked refinement, generating a 4.6 Å map. After CTF correction, Bayesian polishing, a final masked refinement resulted in a 3.93 Å map of DNA-free ComM hexamers (EMDB 19579 PDB 8RXT - Supplementary Figure 9).

### RadA production and purification for biochemical assays

The plasmid IBA13 (Amp^R^, IBA Lifesciences) was previously modified to harbour the gene encoding the superfolder Green Fluorescent Protein (sp-GFP) in frame with the Strep-tag®II encoding-sequence followed by a TEV cleavage site under the Tet promoter. The gene *radA* was cloned from a pneumococcal R800 derivative strain into the modified IBA13 plasmid by FastCloning (Li et al., 2011), downstream of the TEV-cleavage site. This plasmid allows the production of the GFP-RadA fusion protein with a Strep-tag®II at the N-terminus and a TEV cleavage site for the removal of the GFP and the tag.

The RadA-R301A mutant was generated by site-directed mutagenesis PCR on the plasmid using a single primer (EVo-21, see sequence in the R301A mutation in *Streptococcus pneumoniae* part), the PfuTurbo DNA polymerase (Agilent Technologies) and the Taq DNA Ligase (New England Biolabs) according to a protocol adapted from (Shenoy & Visweswariah, 2003).

*E. coli* BL21-Rosetta™(DE3) competent cells (Cm^R^, Novagen) were transformed with the expression vector by heat shock. Transformants were grown at 37 °C in Terrific Broth medium complemented with 100 µg/mL Ampicillin (Amp) and 10 µg/mL Chloramphenicol (Cm) to an optical density of about 0.6. The culture was cooled down for 1h at 16 °C, then the protein expression was induced by adding 200 ng/mL Anhydrotetracycline and incubating at 16 °C overnight. Cells were harvested by centrifugation and resuspended in Sucrose Buffer (25 mM Tris pH 8.0, 25 % sucrose, 1 mM EDTA) complemented with protease inhibitor (cOmplete™, EDTA-free, Sigma-Aldrich) and 200 µg/mL lysozyme and flash-frozen in liquid nitrogen to be stored at −80 °C. Cells were thawed at room temperature and lysed by sonication. Cell debris were pelleted and clear lysate was loaded on Strep-Tactin® resin (IBA Lifesciences) pre-equilibrated with Strep Buffer (50 mM HEPES pH 8.0, 300 mM NaCl, 2 mM MgCl_2_). After a wash step with Strep Buffer, GFP-RadA was eluted with Strep Buffer complemented with 2.5 mM desthiobiotin (IBA Lifesciences). The fusion protein was cleaved with ∼ 80 µg/mL TEV protease (home-made) at 10 °C overnight. Cleaved GFP and desthiobiotin were removed by anion exchange chromatography on a HiTrap Q HP column (GE Healthcare) connected to a FPLC system (ÄKTA purifier-10, GE Healthcare). The cleaved sample was diluted to 100 mM NaCl and loaded on the column pre-equilibrated with Buffer A (50 mM HEPES pH 8.0, 100 mM NaCl, 2 mM MgCl_2_). The column was washed with Buffer A and the proteins were eluted with a linear gradient of 0.1 to 1 M NaCl. RadA was then separated from uncleaved GFP-RadA on StrepTactin® and concentrated by anion exchange chromatography. The final fractions of RadA (containing about 300 mM NaCl) were stored at −20 °C after addition of 50 % glycerol or at −80 °C after addition of 10 % glycerol and freezing in liquid nitrogen.

### DNA substrates for DNA-binding, ATPase and helicase assays

Labeled and unlabeled oligonucleotides were purchased from Eurogentec and Eurofins Genomics respectively.

Single-stranded DNA (ssDNA) for DNA-binding assays is a 100 nt-long oligonucleotide labeled at its 3’ end with the Cy3 dye (Ovio9, see sequence in the table below). Double-stranded DNA (dsDNA) for DNA-binding assays is the 3’Cy3-labeled 100 bp oligonucleotide hybridized to a fully complementary unlabeled 100 nt oligonucleotide (Olea41).

ssDNA for ATPase assays is a 60 nt-long oligonucleotide (Ovio45, see sequence in the table below).

The hybrid DNA substrate for helicase assays is a 30 nt-long oligonucleotide labeled at its 5’ end with the Alexa Fluor 532 (AF532) dye (OA532nt30, see sequence in the table below) hybridized to a 100 nt-long oligonucleotide labeled at its 3’ end with the Black Hole Quencher™ BHQ-1 (oligoBHQ100).

Hybrid and dsDNA substrates for helicase and DNA-binding assays respectively were generated by mixing the two strands at 5 µM each (except oligoBHQ100 at 6 µM, slight excess) in 20 mM Tris pH 7.5, 100 NaCl, 10 mM MgCl_2_, 1 mM dithiothreitol (DTT), incubating 5 min at 100 °C, then placing in boiling water and allowed to cool down to room temperature.

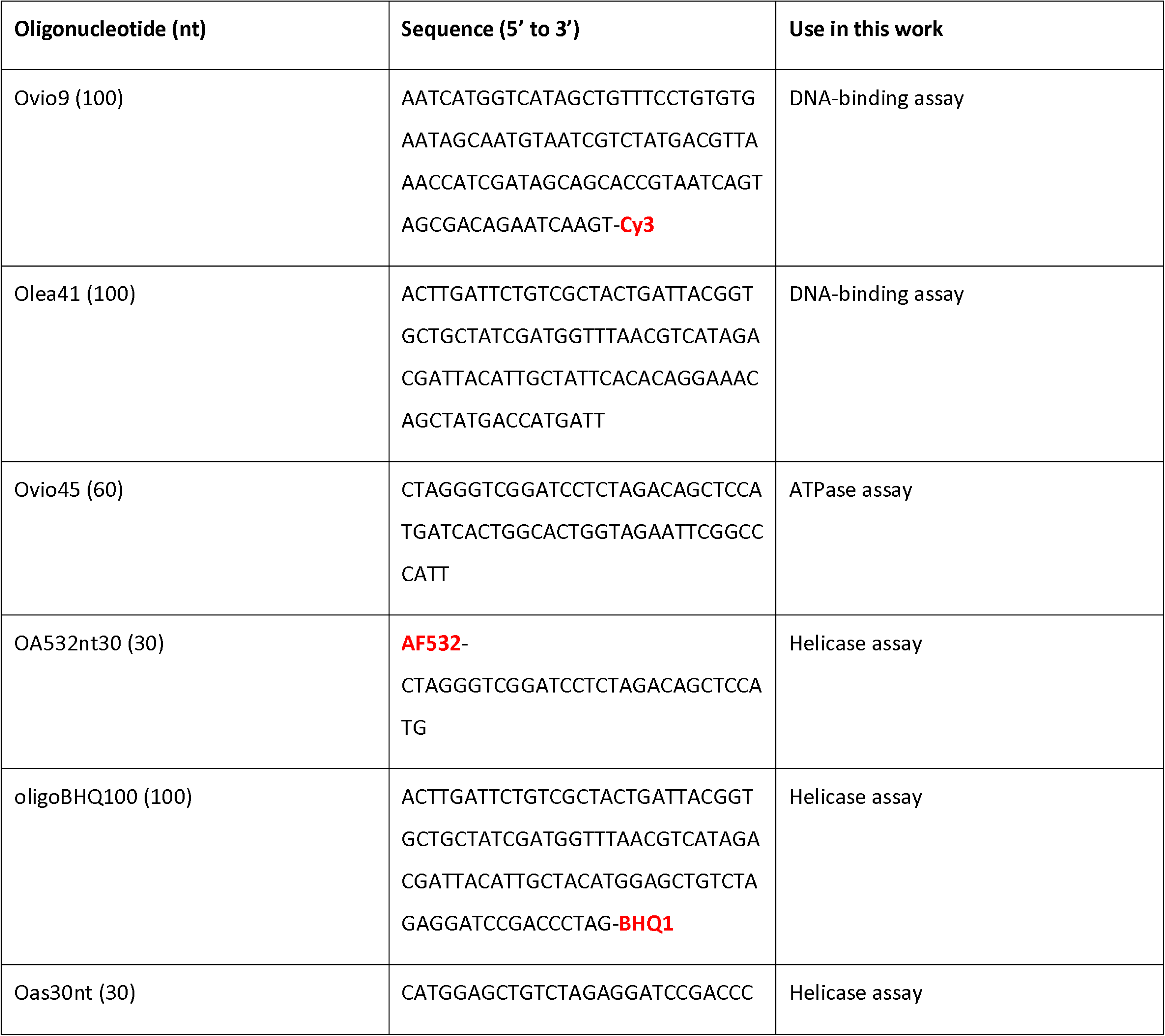
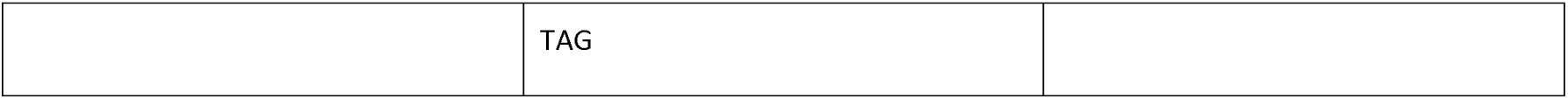

### DNA-binding assays

100, 300 and 900 nM RadA was incubated at room temperature for 15 to 20 min with 10 nM DNA, 10 mM magnesium acetate and 6 % glycerol in a 10 µL reaction solution containing 50 mM Tris pH 7.5, 100 M NaCl, 1 mM DTT, 0.1 mg/mL bovine serum albumin (BSA). Samples were loaded on a 4 % acrylamide gel (acrylamide:bisacrylamide 29:1, 0.3 X TBE) and migrated in 0.3 X TBE for 30 to 70 min at 200 V. The gel was visualized using a Typhoon 9400 (GE Healthcare) with excitation/emission wavelengths of 532/580 nm. The intensity of free DNA bands was quantified using MultiGauge V3.0 (Fujifilm).

### ATPase assays

The ATPase activity of RadA was determined using an ATP regeneration system coupled to NADH oxidation (Kiianitsa et al., 2003; Sehgal et al., 2016). 300 nM RadA was mixed with 1.2 mM phospho(enol)pyruvic acid (Sigma-Aldrich), 70 to 120 U/mL pyruvate kinase (Sigma-Aldrich), 200 U/mL L-lactic dehydrogenase (Sigma-Aldrich), 0.25 mg/mL NADH (Sigma-Aldrich), 2 µM_nt_ ssDNA (Ovio 45) and 2 mM ATP (GE Healthcare) in a 40 µL reaction solution containing 10 mM Tris-HCl pH 7.5, 4 mM magnesium acetate, 0.1 mM DTT. The reaction was carried out in a 96-well plate at 25, 37 or 42 °C on a VarioSkan Flash microplate reader (Thermo Scientific). The NADH absorbance was measured at 340 nm and data were analyzed using Excel (Microsoft). The absorbance decrease was converted to the amount of NADH oxidized (equal to the amount of ATP hydrolyzed) using the NADH extinction coefficient at 340 nm of 6220 M^-1^.cm^-1^.

### Helicase assays

The helicase activity of RadA was determined by measuring the fluorescence intensity increase resulting from the DNA substrate unwinding (separation of fluorescent OA532nt30 and quencher oligoBHQ100 strands). 300 nM RadA was mixed with 10 nM DNA substrate, 100 nM capture strand (Oas30nt, fully complementary to the fluorescent strand to avoid it re-hybridizing to the quencher strand) and 5 mM ATP (GE Healthcare) in a 100 µL reaction solution containing 10 mM Tris pH 7.5, 50 mM NaCl, 10 mM magnesium acetate, 1 mM DTT, 0.1 mg/mL BSA. The reaction was carried out in a quartz cuvette on a FluoroMax spectrofluorimeter (HORIBA) equipped with a circulating water bath set to 37 °C. The fluorescence was measured at 532/554 nm (excitation/emission).

Pneumococcal radA mutagenesis

To generate strain R4735 (with the R301A mutation in RadA), PCR fragments of regions upstream and downstream of the *radA* gene were amplified, with 2 nt of *radA* mutated using primer pairs oALS20-oALS30 and Evo-21-oALS34. Splicing overlap extension (SOE) PCR with these two fragments as templates and the primer pair oALS20-oALS34 was carried out to generate a PCR fragment with the *radA* gene mutated and ∼2,5 kb of homologous sequence on either side. This DNA fragment was transformed into strain R1818 (WT,(Caymaris et al., 2010) without selection. Briefly, 100 μL aliquots of pre-competent cells were resuspended in 900 μL fresh C+Y medium (Martin et al., 1995) (with 100 ng mL^−1^ competence stimulating peptide (CSP) and incubated at 37°C for 10 min. Transforming DNA was then added to a 100 μL aliquot of this culture, followed by incubation at 30°C for 20 min. Cells were then diluted in 1,4 mL C+Y medium and incubated at 37 °C for 3h 30 min, before dilution and plating without selection. Transformants were identified by sequencing.

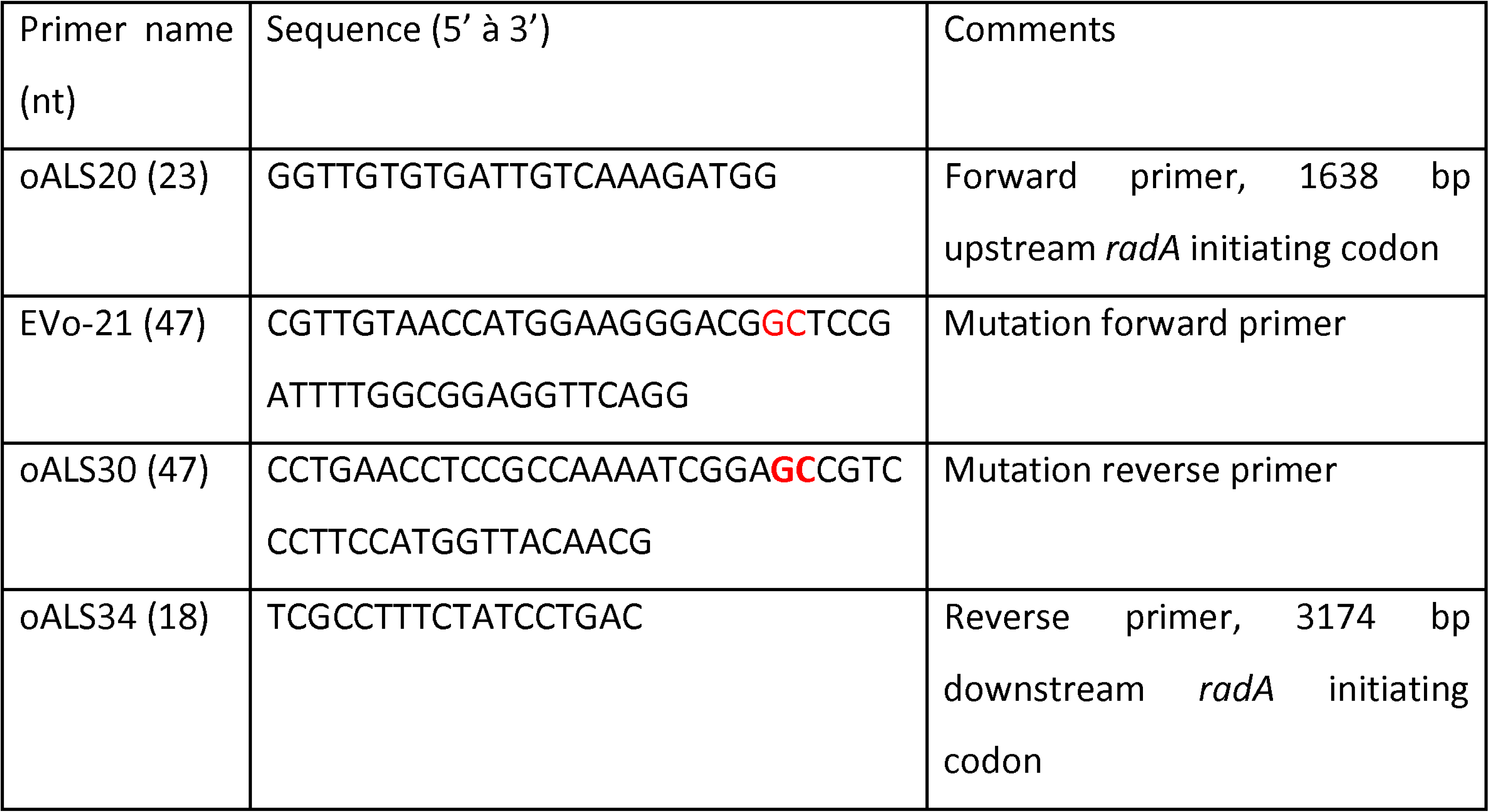

### Transformation assays

To test the efficiency of transformation in wildtype (WT) and mutant strains, 100 µL pre-competent cultures were resuspended in 900 µL fresh C+Y medium pH 7.6, and 100 ng mL CSP was added. Cells were incubated for 10 min at 37°C, before 100 µL was added to tubes containing desired transforming DNA. Transforming DNA was a 4.2 kb-long fragment carrying the rpsL41 Sm allele amplified from R304 chromosomal DNA using the RpsL5-RpsL6 primer pair (Bergé et al., 2013) and providing resistance to Streptomycin. Transforming cultures were incubated at 30 °C for 20 minutes before dilution and plating of appropriate dilutions in 10 mL CAT agar medium (Porter & Guild, 1976) with 3% horse blood. Plates were incubated 2 h at 37°C, before addition of a second 10 mL layer of CAT agar medium with or without Streptomycin for transforming and total cells respectively. Plates were incubated overnight at 37°C and colonies present on selective and non-selective plates were compared to calculate transformation efficiency.

## Data availability

RadA model information is available under accession PDB 8RXC and density map under accession EMDB 19573. dsDNA-bound ComM main map is available under EMDB 19574 and PDB 8RXD, while map relating to subpopulation 2 is under EMDB 19575. Coordinates of ComM Lon domain are available under the accession codes PDB: 8RXS and EMDB 19578. Coordinates for dsDNA-bound ComM trimer are available under accession code PDB: 8RXK and EMDB 19577. Density map for the dsDNA-bound ComM hexamer used for focused refinement are submitted under coordinates EMDB 19581. Coordinates and density map for ComM hexamer in absence of DNA are available under accession codes PDB: 8RXT and EMDB 19579. Density map for ComM dodecamers coordinating dsDNA area available under EMDB 19580 accession code.

## Acknowledgements

This project was executed with funds from the European Research Council Consolidator grant TransfoPneumo. We acknowledge the European Synchrotron Radiation Facility for provision of beam time on CM01 for collection on RadA and we would like to thank Michael Hons for assistance. This work benefited from access to the CryoEM platform of EMBL Heidelberg and has been supported by iNEXT-Discovery, project number 871037, funded by the Horizon 2020 program of the European Commission, with assistance of Felix Weis. EV received a fellowship from the Fondation pour la Recherche Médicale (grant FRM SPF20170938718). We thank Xavier Charpentier for providing the genomic DNA of *L. pneumophila paris*.

---

Supplementary table 1: Synthetic oligonucleotides used for in vitro reconstitution of helicase hexamers

Supplementary table 2: PDB-PISA analysis of RadA and ComM Lon domain hexamers

Supplementary Table 3: CryoEM collect and modeling statistics

Supplementary Figure 1: **CryoEM data collection, processing and modeling of RadA.**

(a) Representative micrograph for the collect on RadA. The red circle shows a typical top view of RadA, while the white circle represents a side view.

(b) representative 2D classes after particle classification in Relion 3.1.

(c) Particle orientation distribution, plotted on the Euler sphere around a representation of the RadA CryoEM map.

(d) Fourrier Shell Correlation (FSC) map obtained from half-maps in Phenix Autosharpen software.

(e) Local-filtered map of RadA at level 0.015, colored by local resolution (in Å), with a slice of the central region of RadA (right panel).

(f) Representative regions of the CryoEM map depicted as a blue mesh, with respective model fit to the density. Chain identifier and residues number for the modelled regions are shown above each image.

Supplementary Figure 2: **Comparison between RadA and the ATPase region of the Superfamily 4 helicase T4-gp4 (PDB: 6N7I)**

(a) Overall structure of RadA (top) and T7-gp4 (middle). Both helicases present a helical arrangement, tilted in relation to the DNA helix. Bottom panel depicts RadA in ribbon representation, fit inside the transparent surface density of T7-gp4.

(b) Top view from the C-terminal of RadA (top panel) and T7-gp4 (bottom panel)

(c) ATP coordination sites of RadA (top) and T7-gp4 (bottom). ATP-y-S (in RadA) and ATP (in T7-gp4) are depicted as sticks colored by element. Ribbon representation for Chain D is shown in green and chain E in cyan. Relevant residues are shown as sticks, colored by element.

(d) Cooperative coordination of ATP and DNA in RadA (top) and T7-gp4 (bottom). ATP-y-S (in RadA) and ATP (in T7-gp4), as well as relevant residues, are depicted in stick representation colored by elements. The ATPase domains are depicted as green ribbon, with the region structured upon ATP binding highlighted in golden. DNA is depicted as sticks, with the bases represented as blue rectangles.

(e) Superposition of the ATPase domains of RadA (orange, residues 54 to 273) and T7-gp4 (blue, residues 263 to 547), resulting in an RMSD of 1.16 Å over 120 pruned residues or 1.944 Å overall.

(f) ConSurf (REF) analysis of the ATPase domain of RadA, showing conservation of residues along homologous sequences, depicted in a color gradient from cyan (variable) to purple (conserved).

Supplementary Figure 3: **CryoEM data collection, processing and modeling of ComM hexamer bound to DNA.**

(a) Representative micrograph for the collect on ComM. The red circle shows a typical top view of ComM, while the white circle represents a side view.

(b) representative 2D classes after particle classification in Cryosparc.

(c) Particle orientation distribution of ComM focused map on DNA-bound ComM hexamer. (d) Fourrier Shell Correlation (FSC) map obtained from half-maps in Cryosparc.

(e) Local-filtered map of ComM hexamer, at level 0.2, colored by local resolution (in Å). Right panel shows a cross-section of the map.

(f) Representative regions of the CryoEM map depicted as a blue mesh, with respective model fit to the density. Chain identifier and residues number for the modeled regions are shown above each image.

Supplementary Figure 4: **local refinement of DNA-bound ComM hexamer.**

Local refinement on chains C-E + DNA (a to c).

(a) Particle orientation distribution of ComM focused map on chains C-E + DNA, obtained in Cryosparc.

(b) Fourrier Shell Correlation (FSC) map obtained from half-maps in Cryosparc.

(c) Local-filtered map of ComM chains C-E + DNA, at level 0.07, coloured by local resolution (in Å).

Local refinement on the Lon domain (d to f).

(d) Particle orientation distribution of ComM focused map on Lon domains. (e) Fourrier Shell Correlation (FSC) map obtained from half-maps in Cryosparc.

(f) Local-filtered map of ComM Lon domains, at level 0.07, coloured by local resolution (in Å). Local refinement on the subpopulation 2 DNA-bound ComM hexamer (g to i).

(g) Particle orientation distribution of ComM focused map on Lon domains. (h) Fourrier Shell Correlation (FSC) map obtained from half-maps in Cryosparc.

(i) Local-filtered map of ComM Lon domains, at level 0.07, coloured by local resolution (in Å).

Supplementary Figure 5: **Comparison between ComM and the C-terminal ATPase domains of human MCM (PDB: 6SKL), RavA (PDB 6SZB) and Magnesium Chelatase (PDB 1G8P).**

(a) Arrangement around the DNA for ComM (top) and MCM (bottom), with MCM colored according to ComM scheme, from MCM7 to MCM4.

(b) View from the top on the N-terminal side of ComM (top) and MCM (bottom).

(c) ATP coordination site in ComM (top) and MCM (bottom). In ComM, AMP-PNP is depicted in sticks, colored by element, with the cryoEM density map depicted as a blue mesh. Chains C and D are depicted as yellow and green ribbons, respectively, with relevant residues depicted in sticks and coloured by element. In MCM, MCM2 is depicted in green, while MCM6 is depicted in cyan. AMP-PNP is depicted as sticks, coloured by element, and so are relevant residues.

(d) Comparison between DNA binding loops of ATPase domains in ComM (top) and MCM (bottom). The ATPase domains of ComM monomer D and MCM2 are depicted in green ribbons, with ps1B loop depicted in pink and H2i in golden.

(e) Superposition of ComM, RavA and Mg^2+^-chelatase. ComM ATPase domain (residues 190-500) is shown in orange, RavA ATPase domain is depicted in blue (PDB 6SZB, residues 3-306), and shows an RMSD of 2.176 Å with ComM. Mg^2+^ Chelatase (PDB 1G8P) is shown in green, and has an RMSD of 1.928 Å with ComM.

(f) Consurf analysis of ComM ATPase domain (residues 190 to 500), showing conservation of residues along homologous sequences, depicted in a color gradient from cyan (variable) to purple (conserved).

Supplementary Figure 6: **CryoEM data collection, processing and modeling of DNA-free ComM hexamer.**

(a) Particle orientation distribution of DNA-free ComM hexamer, plotted in the Euler-angle sphere around the density map.

(b) Fourrier Shell Correlation (FSC) map obtained from half-maps in Relion 4.0.

(c) Local-filtered map of DNA-free ComM hexamer, at level 0.002, coloured by local resolution(in Å).

(d) CryoEM analysis of DNA-free ComM hexamer. Despite unchanged position for the Lon domains (not shown), the ATPase domains are organized as a pair of trimers, showing C2 symmetry. Four molecules of AMP-PNP are orchestrated in the internal interfaces of each trimer (A-B, B-C, D-E, E-F), but not in the limit between the trimers (A-F and C-D). The 4 Å density map is despicted at level 0.002 (upper panel), coloured accordingly to the molecular model (lower panel)

Supplementary Figure 7: **CryoEM data collection, processing and modeling of ComM dodecamer.**

(a) Particle orientation distribution of ComM dodecamer.

(b) Fourrier Shell Correlation (FSC) map obtained from half-maps in Cryosparc.

(c) Local-filtered map of ComM dodecamer, at level 0.04, coloured by local resolution(in Å). (d) CryoEM density of ComM dodecamers, shown with 70% transparency at level 0.04. Two ComM hexamers are fitted in the density, shown in blue and green ribbons, alongside a dsDNA molecule in magenta and golden encompassing the two hexamers. In between the fitted hexamers, an additional density is observed in the position equivalent to the expected ZF domain, highlighted in orange.

(e) Density corresponding to the 47 bp dsDNA encompassing both hexamers in the ComM dodecamer. Density for the DNA is shown at 50 % transparency while the rest of the density is hidden. Fitted dsDNA is depicted as ribbons in magenta and golden, with bases represented as sticks and colored by heteroatom. ComM hexamers are shown as blue and green ribbons, with 70% transparency.

Supplementary Figure 8: **Comparison between two ComM hexamer populations**.

On subpopulation 1 (left panel – EMDB 19574), A and F subunits (red and blue, respectively) are disengaged in relation to each other. In subpopulation 2 (central panel – EMDB 19575), A and F subunits are closely engaged, with consequent disengagement of F from E (cyan). The right panel shows superposition of subunits A and F, aligned on the Lon domains, for subpopulation 1 (gray ribbons) and subpopulation 2 (coloured ribbons).

Supplementary Figure 9: **CryoEM processing workflow for RadA and ComM**

Supplementary video 1**: 360 degree rotation of RadA, depicted as transparent ribbons, with only the DNA binding loops colored in the color-scheme of figure 2**. DNA backbone is shown in ribbon representation, with magenta color for leading strand and golden for lagging strand. Nucleotide bases are depicted as blue rectangles.

Supplementary video 2**: 360 degree rotation of ComM, depicted as transparent ribbons, with only the DNA binding loops colored in the color-scheme of figure 2**. DNA backbone is shown in ribbon representation, with magenta color for leading strand and golden for lagging strand. Nucleotide bases are depicted as blue rectangles.

Supplementary video 3: **Morphing of RadA conformations, by alternating the position of each monomer in sequence**

Supplementary video 4: **Morphing of ComM hexamer subpopulation, evidencing movement in chains A and F.**

Supplementary video 5: **Morphing of ComM monomer conformations, by alternating the position of each monomer in sequence.**

Supplementary video 6: **Morphing of the six different monomers constituting the RadA hexamer (A-B-C-D-E-F-A), aligned on the Lon domains.** The ATPase elongate from the Lon domain along the y axis, before disengaging and retracting again. Additionally, it rotates on its own X axis, in a hinge movement. The salt bridge switch of R301 is highlighted, initially contacting E239, and then changing to contact with E76.

