## Supplementary information for "Structural basis for activation and mechanism of the bacterial hexameric motors RadA and ComM in DNA branch migration during homologous recombination"

**1.** Structure and Function of Bacterial Nanomachines – Institut Européen de Chimie et Biologie, Microbiologie fondamentale et pathogénicité, UMR 5234, CNRS, University of Bordeaux, 2 rue Robert Escarpit, 33600, Pessac, France

**2.** Laboratoire de Microbiologie et de Génétique Moléculaire (UMR 5100). Centre de Biologie Intégrative; 169, avenue Marianne Grunberg-Manago; CNRS - Université Paul Sabatier - 31062 Toulouse cedex 09 France

3. Present address: Departamento de Bioquímica e Biologia Tecidual. Laboratório de Bioquímica de Complexos Bacterianos. Instituto de Biologia. Universidade Estadual de Campinas (UNICAMP)– Rua monteiro Lobato, 255 – Campinas-SP, Brasil 13083-862

4. Present address : TBI, Université de Toulouse, CNRS, INRAE, INSA, Toulouse, France

*correspondance :
### These authors contributed equally to this work.

**Supplementary Table 1:** Synthetic oligonucleotides used for in vitro reconstitution of helicase hexamers.

| Synthetic DNA sequences | |
|---|---|
| A | 20A-60C |
| B | 20T |
| C | 30A-60C-30A |
| D | 30T |
| E | 60A-60C |
| F | 60A |
| G | 60T |
| H | 60G |

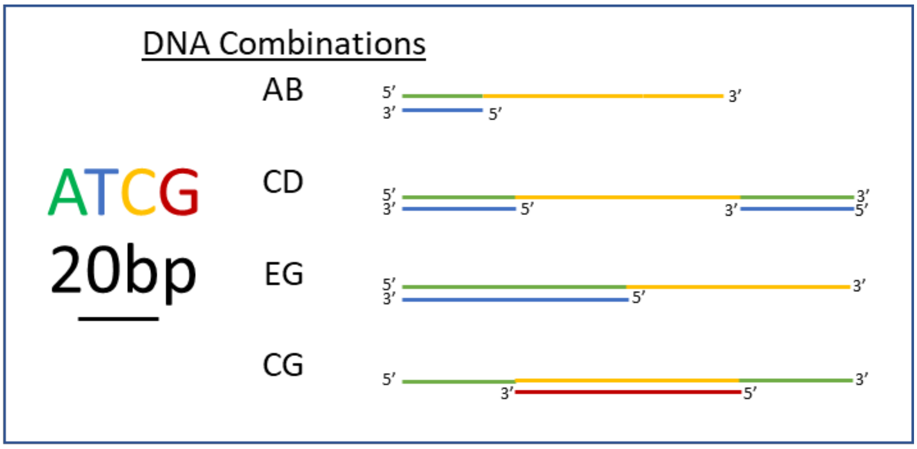

**Supplementary Table 2:** PDB-PISA analysis of RadA and ComM Lon domain hexamers.

| | Solvent acessible surface | Number of interface residues | Interface area | Solvatation free energy ∆G kCal/mol | ∆G p-value |
|---|---|---|---|---|---|
| ComM | 8206 $Å^2$ | 21 | 678 $Å^2$ | -3.9 | 0.477 |
| RadA | 8947 $Å^2$ | 32 | 1028 $A^2$ | -6.8 | 0.28 |

**Supplementary Table 3:** CryoEM collect and modeling statistics.

| Dscription | DNA-bound ComM hexamer-Main Map | DNA-bound ComM hexamer-subpopulation 2 | ComM hexamer consensus | ComM Trimer | ComM Lon domains | DNA-free ComM | ComM Dodecamer | RadA |
|---|---|---|---|---|---|---|---|---|
| PDB | 8RXD | | | 8RXK | 8RXS | 8RXT | | 8RXC |
| EMDB | 19574 | 19575 | 19581 | 19577 | 19578 | 19579 | 19580 | 19573 |
| **Data collection** | | | | | | | | |
| Microscope | Titan Krios | | | | | | | Titan Krios |
| camera | Quantum K3 | | | | | | | Falcon 3 |
| Voltage (kV) | 300 | | | | | | | 300 |
| Magnification | 130,000 | | | | | | | 165 k |
| Electron exposure (e– per A▯) | 1.339 | | | | | | | 1.2 |
| Total Dose | 53.56 | | | | | | | 54.05 |
| Pixel size (A▯ | 0.645 | | | | | | | 0.827 |
| Decofus range (um) | -0.6 to -1.8 | | | | | | | -0.5 to -2.5 |
| **Processing** | | | | | | | | |
| Symmetry imposed | C1 | | | | | | | C1 |
| Micrographs number | 32,534 | | | | | | | 8983 |
| Initial particle images (no.) | | | 1,313,661 | | | 1,249,471 | 1,313,661 | 3,768,530 |
| Final particle images (no.) | 207,223 | 105,754 | 852,356 | 305,846 | 852,356 | 78,222 | 69,029 | 97,546 |
| Map resolution (Å)-(0.143 FSC threshold model) | 3.13 | 3.37 | 3 | 3.23 | 2.8 | 3.93 | 4.1 | 3.15 |
| **Refinement and validation** | | | | | | | | |
| Map sharpening (B-factor) (A▯2) | 80.6 | 75.8 | 117.12 | 103.1 | 92 | 82 | 58 | 101 |
| **Model composition** | | | | | | | | |
| No. of chains | 12 | | | 9 | 6 | 10 | | 14 |
| Atoms (no.) | 23172 | | | 12162 | 16267 | 22258 | | 19206 |
| Aminoacid Residues (no.) | 2898 | | | 1446 | 1080 | 2891 | | 2376 |
| Nucleotide Residues | 48 | | | 48 | 0 | 0 | | 42 |
| Ligands (no.) | 4 | | | 3 | 0 | 4 | | 10 |
| Bond lengths (A▯ | 0.005 | | | 0.003 | 0.003 | 0.004 | | 0.004 |
| Bond angles (°) | 0.734 | | | 0.653 | 0.609 | 0.846 | | 0.713 |
| Ramachandran favored % | 90.86 | | | 92.17 | 93.07 | 93.01 | | 94.75 |
| Ramachandran allowed % | 8.26 | | | 7.13 | 6.65 | 6.33 | | 4.74 |
| Ramachandran outliers % | 0.87 | | | 0.7 | 0.28 | 0.66 | | 0.51 |
| Rotamers outliers % | 0.75 | | | 0.25 | 0.45 | 0.62 | | 0.25 |
| MolProbity score | 2.08 | | | 1.97 | 1.93 | 1.89 | | 1.82 |
| Clashscore | 10.8 | | | 9.26 | 9.16 | 8.22 | | 8.52 |
| CC (mask) | 0.83 | | | 0.73 | 0.62 | 0.78 | | 0.25 |
| CC (box) | 0.39 | | | 0.54 | 0.4 | 0.5 | | 0.34 |
| CC (peaks) | 0.43 | | | 0.39 | 0.28 | 0.46 | | 0.18 |
| CC (volume) | 0.82 | | | 0.73 | 0.62 | 0.76 | | 0.27 |
| Mean CC for Ligands | 0.76 | | | 0.62 | | 0.71 | | 0.23 |

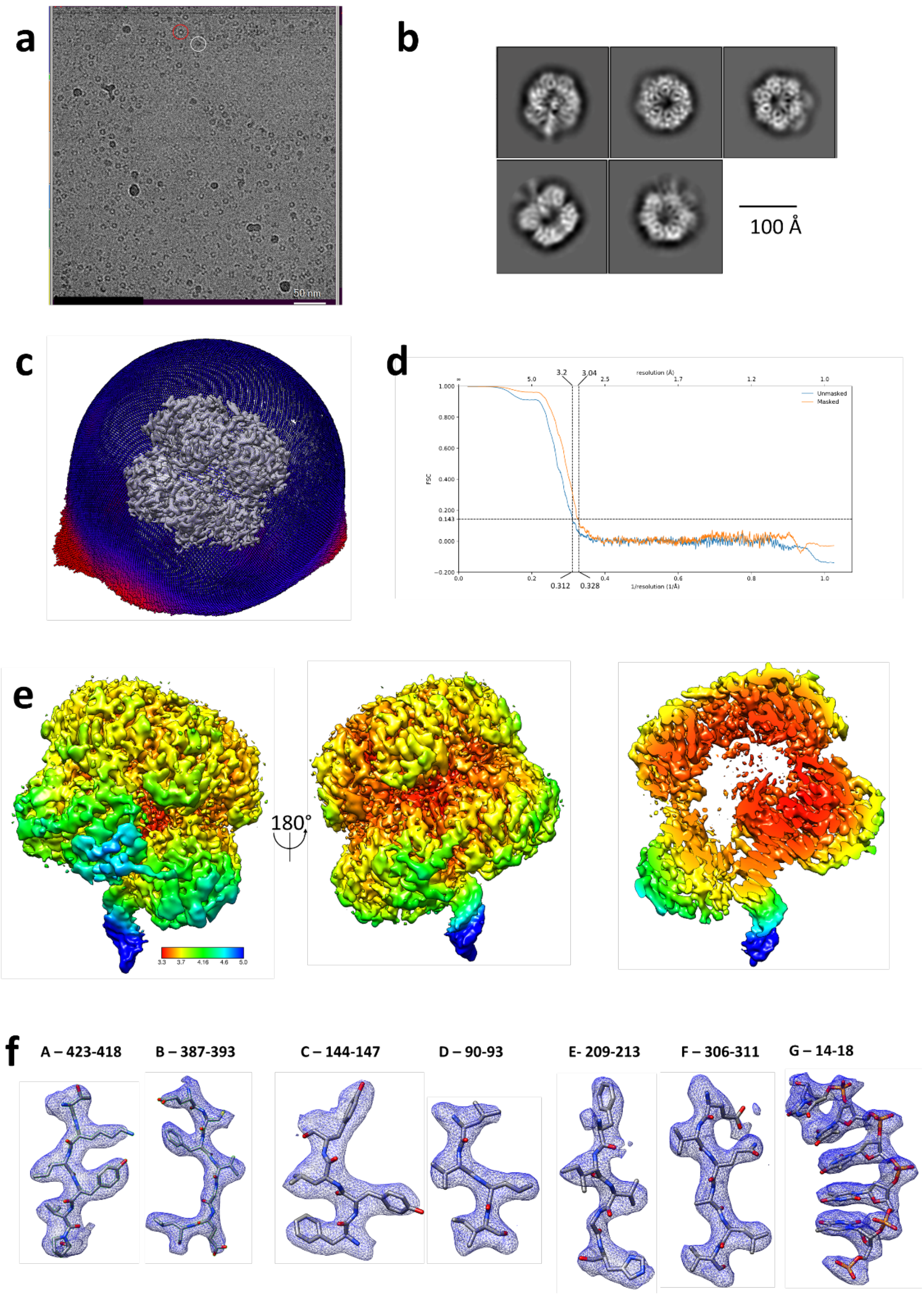

**Supplementary Figure 1: CryoEM data collection, processing and modeling of RadA.** (**a**) Representative micrograph for the collect on RadA. The red circle shows a typical top view of RadA, while the white circle represents a side view. (**b**) representative 2D classes after particle classification in Relion 3.1. (**c**) Particle orientation distribution, plotted on the Euler sphere around a representation of the RadA CryoEM map. (**d**) Fourrier Shell Correlation (FSC) map obtained from half-maps in Phenix Autosharpen software. (**e**) Local-filtered map of RadA at level 0.015, colored by local resolution (in Å), with a slice of the central region of RadA (right panel). (**f**) Representative regions of the CryoEM map depicted as a blue mesh, with respective model fit to the density. Chain identifier and residues number for the modelled regions are shown above each image.

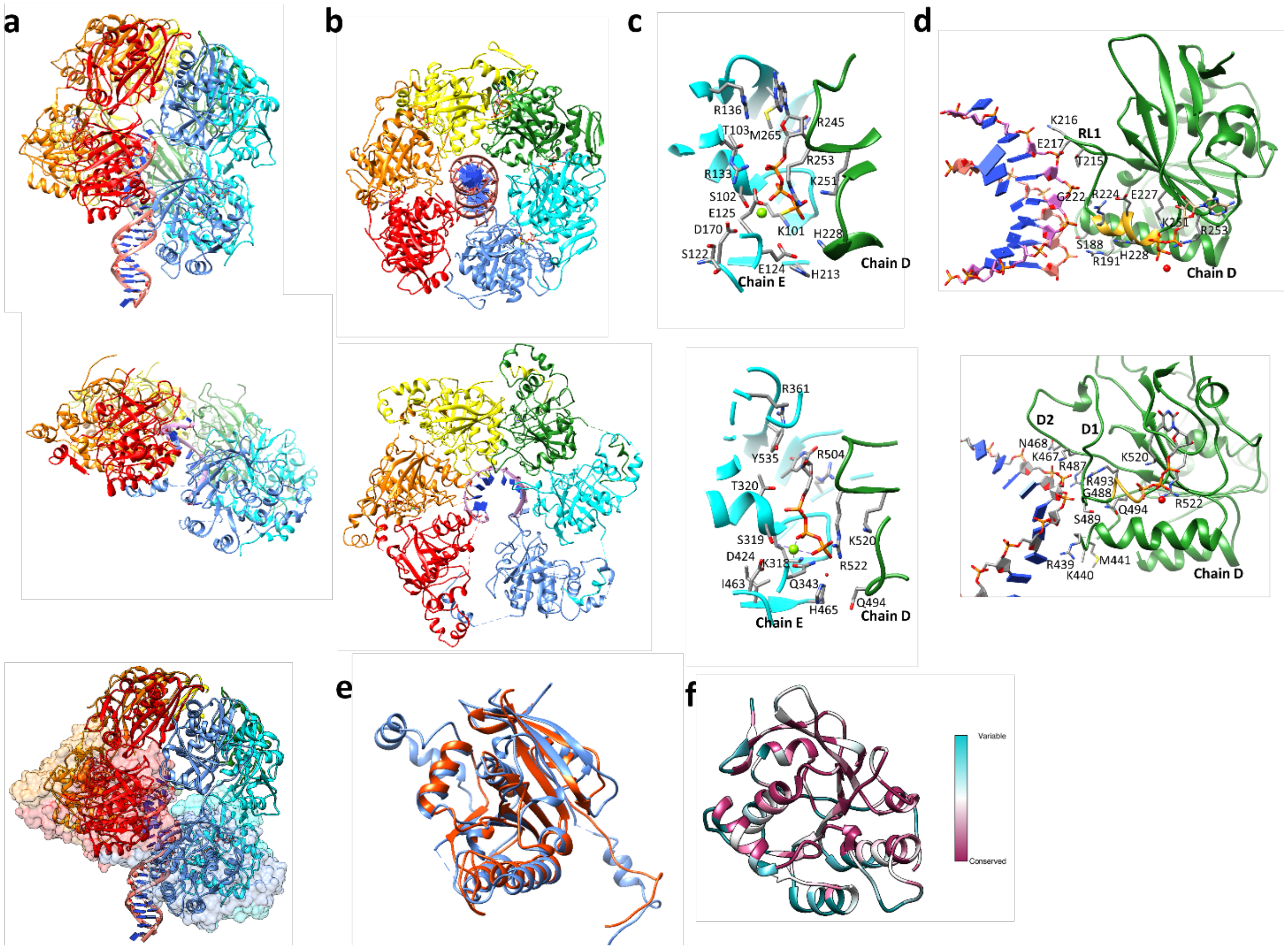

**Supplementary Figure 2: Comparison between RadA and the ATPase region of the Superfamily 4 helicase T4-gp4 (PDB: 6N7I).** (**a**)Overall structure of RadA (top) and T7-gp4 (middle). Both helicases present a helical arrangement, tilted in relation to the DNA helix. Bottom panel depicts RadA in ribbon representation, fit inside the transparent surface density of T7-gp4. (**b**)Top view from the C-terminal of RadA (top panel) and T7-gp4 (bottom panel). (**c**) ATP coordination sites of RadA (top) and T7-gp4 (bottom). ATP-y-S (in RadA) and ATP (in T7-gp4) are depicted as sticks colored by element. Ribbon representation for Chain D is shown in green and chain E in cyan. Relevant residues are shown as sticks, colored by element. (**d**) Cooperative coordination of ATP and DNA in RadA (top) and T7-gp4 (bottom). ATP-y-S (in RadA) and ATP (in T7-gp4), as well as relevant residues, are depicted in stick representation colored by elements. The ATPase domains are depicted as green ribbon, with the region structured upon ATP binding highlighted in golden. DNA is depicted as sticks, with the bases represented as blue rectangles. (**e**) Superposition of the ATPase domains of RadA (orange, residues 54 to 273) and T7-gp4 (blue, residues 263 to 547), resulting in an RMSD of 1.16 Å over 120 pruned residues or 1.944 Å overall. (**f**) ConSurf (REF) analysis of the ATPase domain of RadA, showing conservation of residues along homologous sequences, depicted in a color gradient from cyan (variable) to purple (conserved).

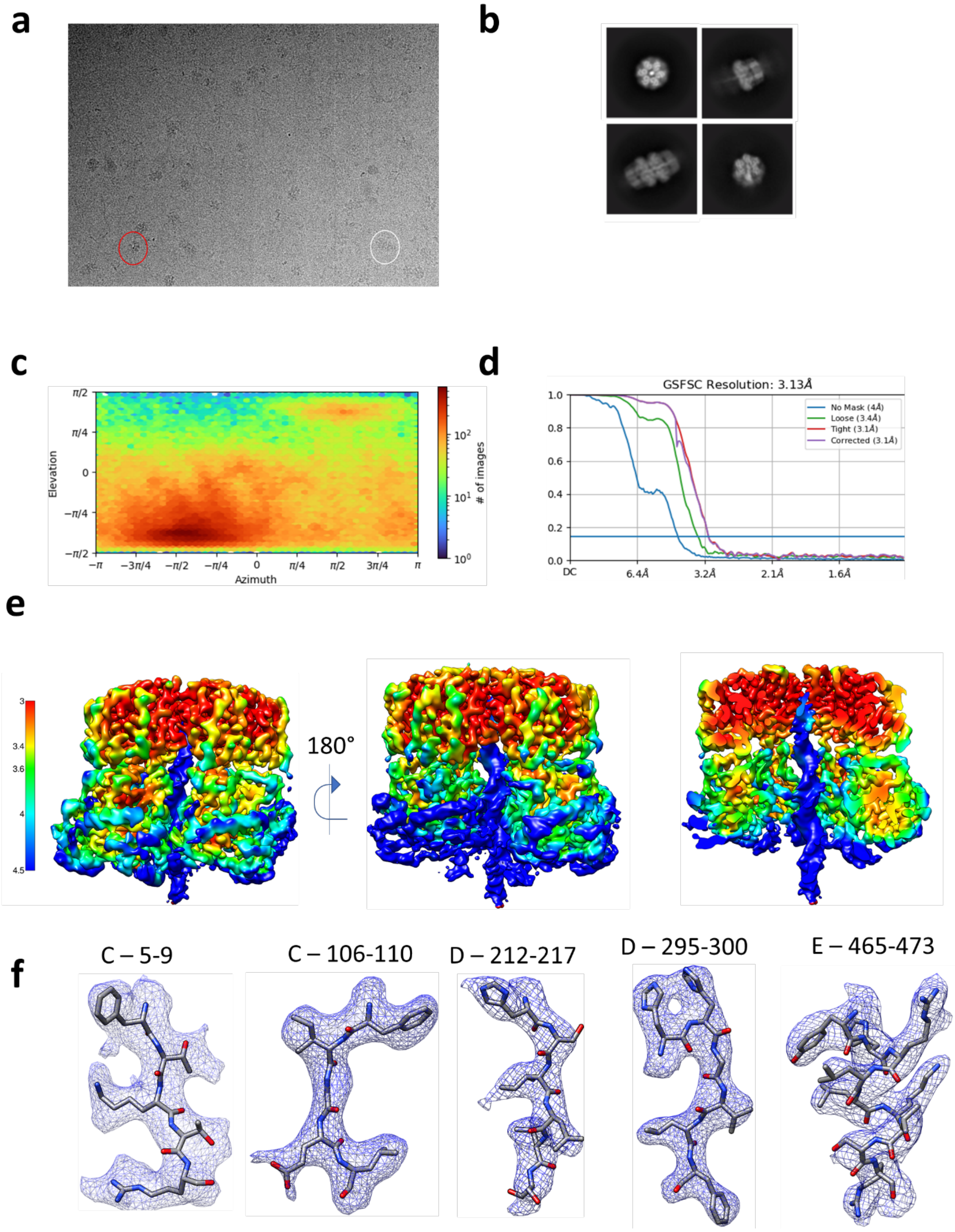

**Supplementary Figure 3: CryoEM data collection, processing and modeling of ComM hexamer bound to DNA.** (**a**) Representative micrograph for the collect on ComM. The red circle shows a typical top view of ComM, while the white circle represents a side view. (**b**) representative 2D classes after particle classification in Cryosparc. (**c**) Particle orientation distribution of ComM focused map on DNA-bound ComM hexamer. (**d**) Fourrier Shell Correlation (FSC) map obtained from half-maps in Cryosparc. (**e**) Local-filtered map of ComM hexamer, at level 0.2, colored by local resolution (in Å). Right panel shows a cross-section of the map. (**f**) Representative regions of the CryoEM map depicted as a blue mesh, with respective model fit to the density. Chain identifier and residues number for the modelled regions are shown above each image.

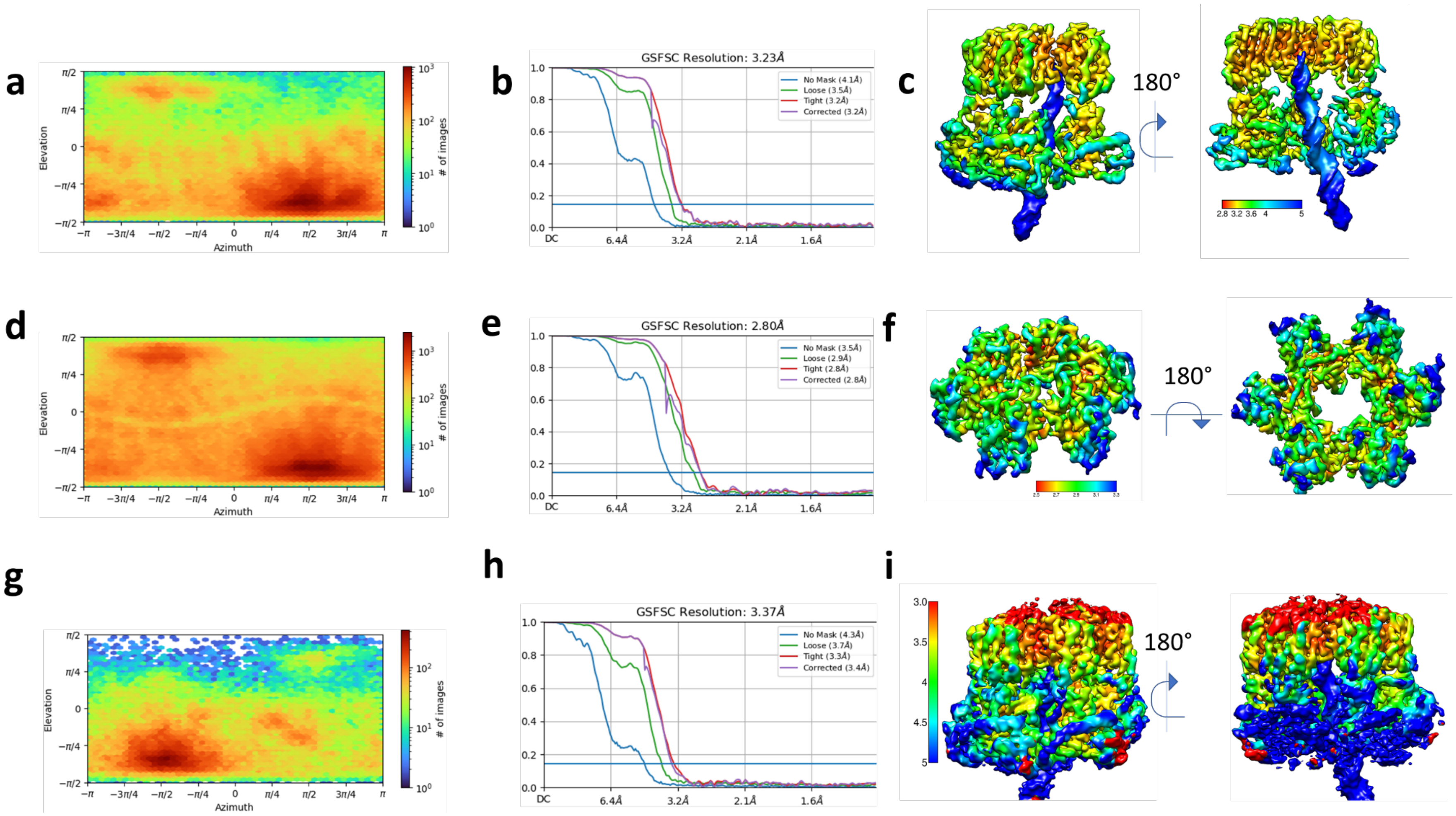

**Supplementary Figure 4: Local refinement of DNA-bound ComM hexamer.** Local refinement on chains C-E + DNA (**a to c**). (**a**) Particle orientation distribution of ComM focused map on chains C-E + DNA, obtained in Cryosparc. (**b**) Fourrier Shell Correlation (FSC) map obtained from half-maps in Cryosparc. (**c**) Local-filtered map of ComM chains C-E + DNA, at level 0.07, coloured by local resolution (in Å). Local refinement on the Lon domain (**d to f**). (**d**) Particle orientation distribution of ComM focused map on Lon domains. (**e**) Fourrier Shell Correlation (FSC) map obtained from half-maps in Cryosparc. (**f**) Local-filtered map of ComM Lon domains, at level 0.07, coloured by local resolution (in Å). Local refinement on the subpopulation 2 DNA-bound ComM hexamer (**g to i**). (**g**) Particle orientation distribution of ComM focused map on Lon domains. (**h**) Fourrier Shell Correlation (FSC) map obtained from half-maps in Cryosparc. (**i**) Local-filtered map of ComM Lon domains, at level 0.07, coloured by local resolution (in Å).

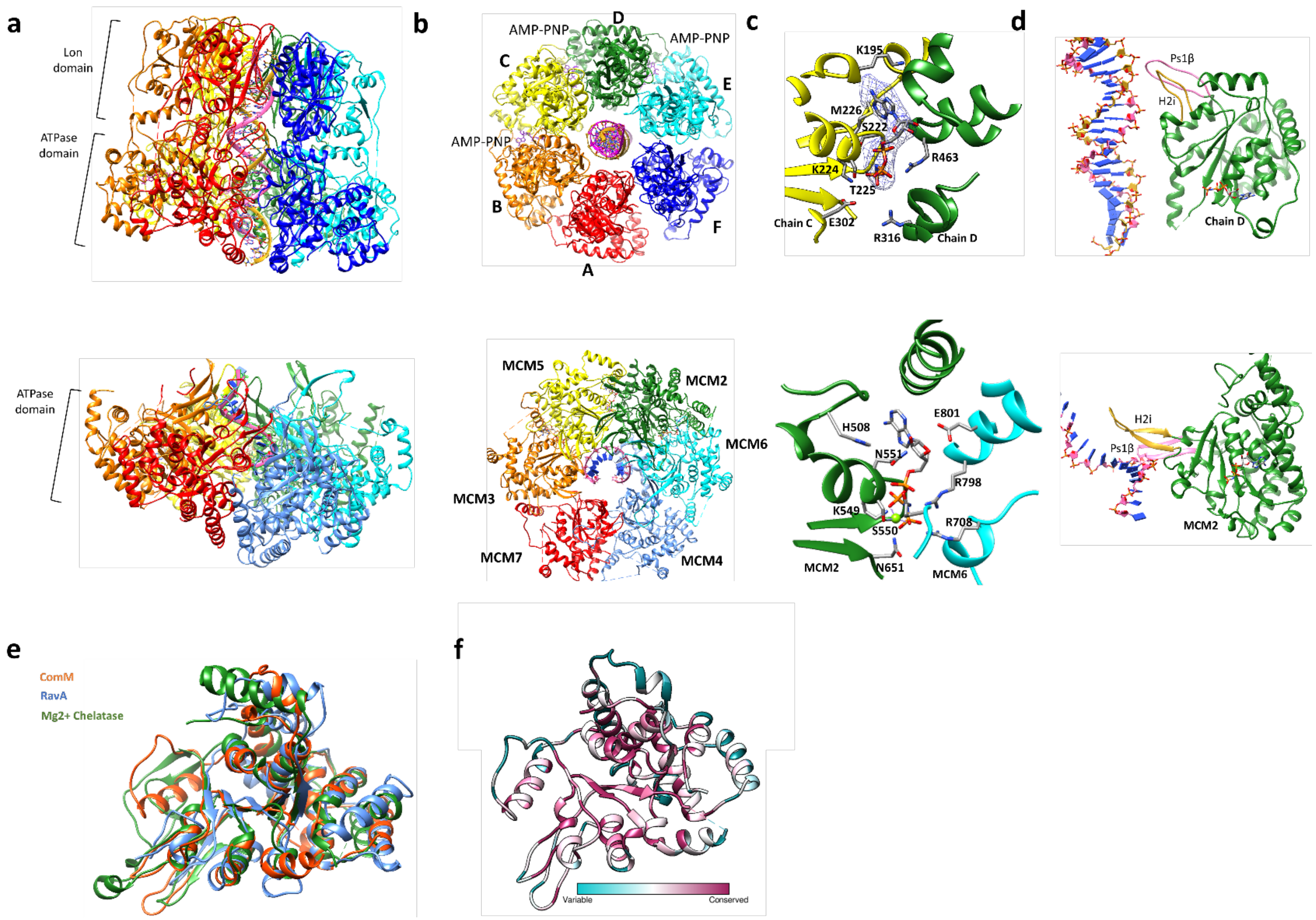

**Supplementary Figure 5: Comparison between ComM and the C-terminal ATPase domains of human MCM (PDB: 6SKL), RavA (PDB 6SZB) and Magnesium Chelatase (PDB 1G8P).** (**a**) Arrangement around the DNA for ComM (top) and MCM (bottom), with MCM colored according to ComM scheme, from MCM7 to MCM4. (**b**) View from the top on the N-terminal side of ComM (top) and MCM (bottom). (**c**) ATP coordination site in ComM (top) and MCM (bottom). In ComM, AMP-PNP is depicted in sticks, colored by element, with the cryoEM density map depicted as a blue mesh. Chains C and D are depicted as yellow and green ribbons, respectively, with relevant residues depicted in sticks and coloured by element. In MCM, MCM2 is depicted in green, while MCM6 is depicted in cyan. AMP-PNP is depicted as sticks, coloured by element, and so are relevant residues. (**d**) Comparison between DNA binding loops of ATPase domains in ComM (top) and MCM (bottom). The ATPase domains of ComM monomer D and MCM2 are depicted in green ribbons, with ps1B loop depicted in pink and H2i in golden. (**e**) Superposition of ComM, RavA and $Mg^{2+}$-chelatase. ComM ATPase domain (residues 190-500) is shown in orange, RavA ATPase domain is depicted in blue (PDB 6SZB, residues 3-306), and shows an RMSD of 2.176 Å with ComM. $Mg^{2+}$ Chelatase (PDB 1G8P) is shown in green, and has an RMSD of 1.928 Å with ComM. (**f**) Consurf analysis of ComM ATPase domain (residues 190 to 500), showing conservation of residues along homologous sequences, depicted in a color gradient from cyan (variable) to purple (conserved).

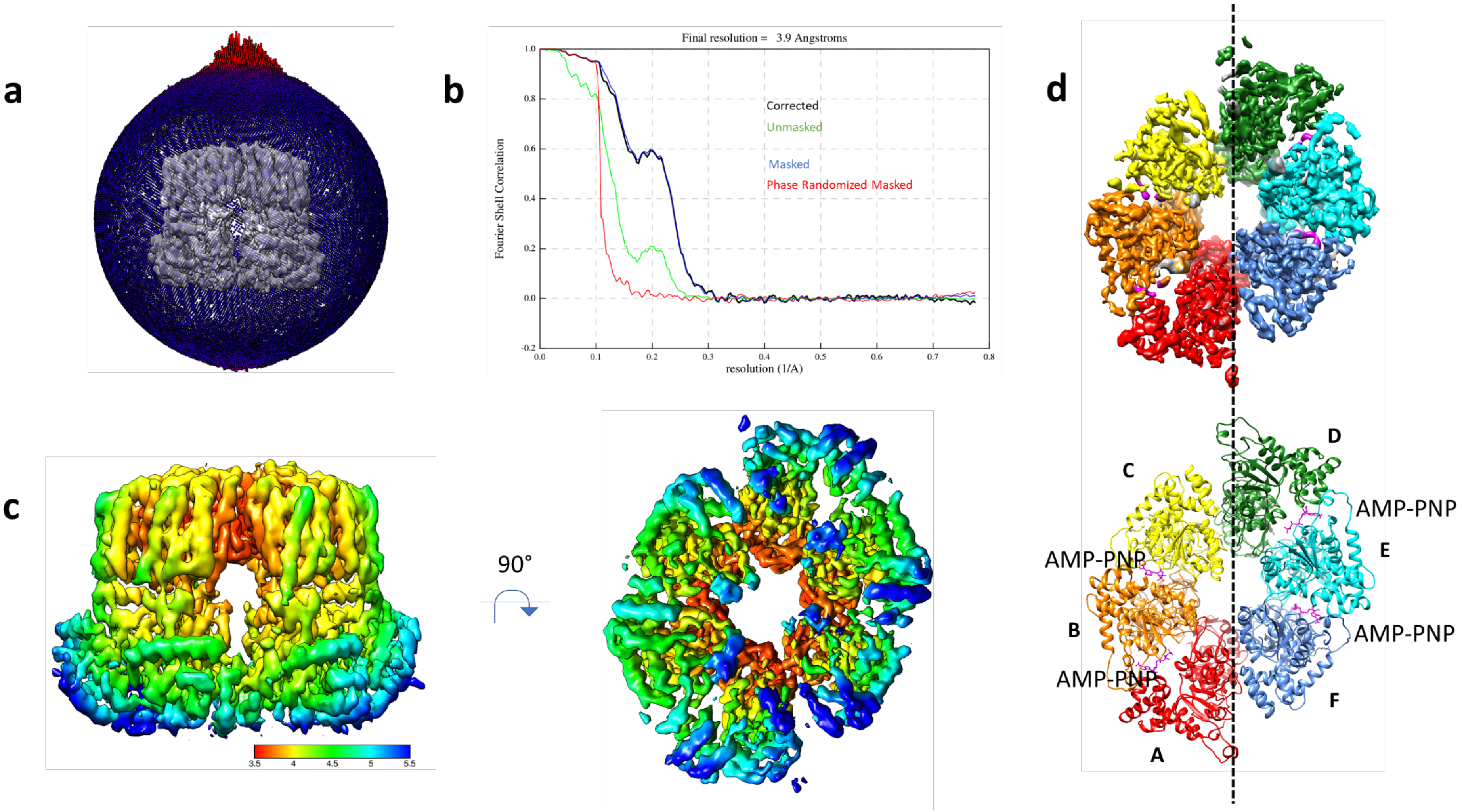

**Supplementary Figure 6: CryoEM data collection, processing and modeling of DNA-free ComM hexamer.** (**a**) Particle orientation distribution of DNA-free ComM hexamer, plotted in the Euler-angle sphere around the density map. (**b**) Fourrier Shell Correlation (FSC) map obtained from half-maps in Relion 4.0. (**c**) Local-filtered map of DNA-free ComM hexamer, at level 0.002, coloured by local resolution (in Å). (**d**) CryoEM analysis of DNA-free ComM hexamer. Despite unchanged position for the Lon domains (not shown), the ATPase domains are organized as a pair of trimers, showing C2 symmetry. Four molecules of AMP-PNP are orchestrated in the internal interfaces of each trimer (A-B, B-C, D-E, E-F), but not in the limit between the trimers (A-F and C-D). The 4 Å density map is despicted at level 0.002 (upper panel), coloured accordingly to the molecular model (lower panel).

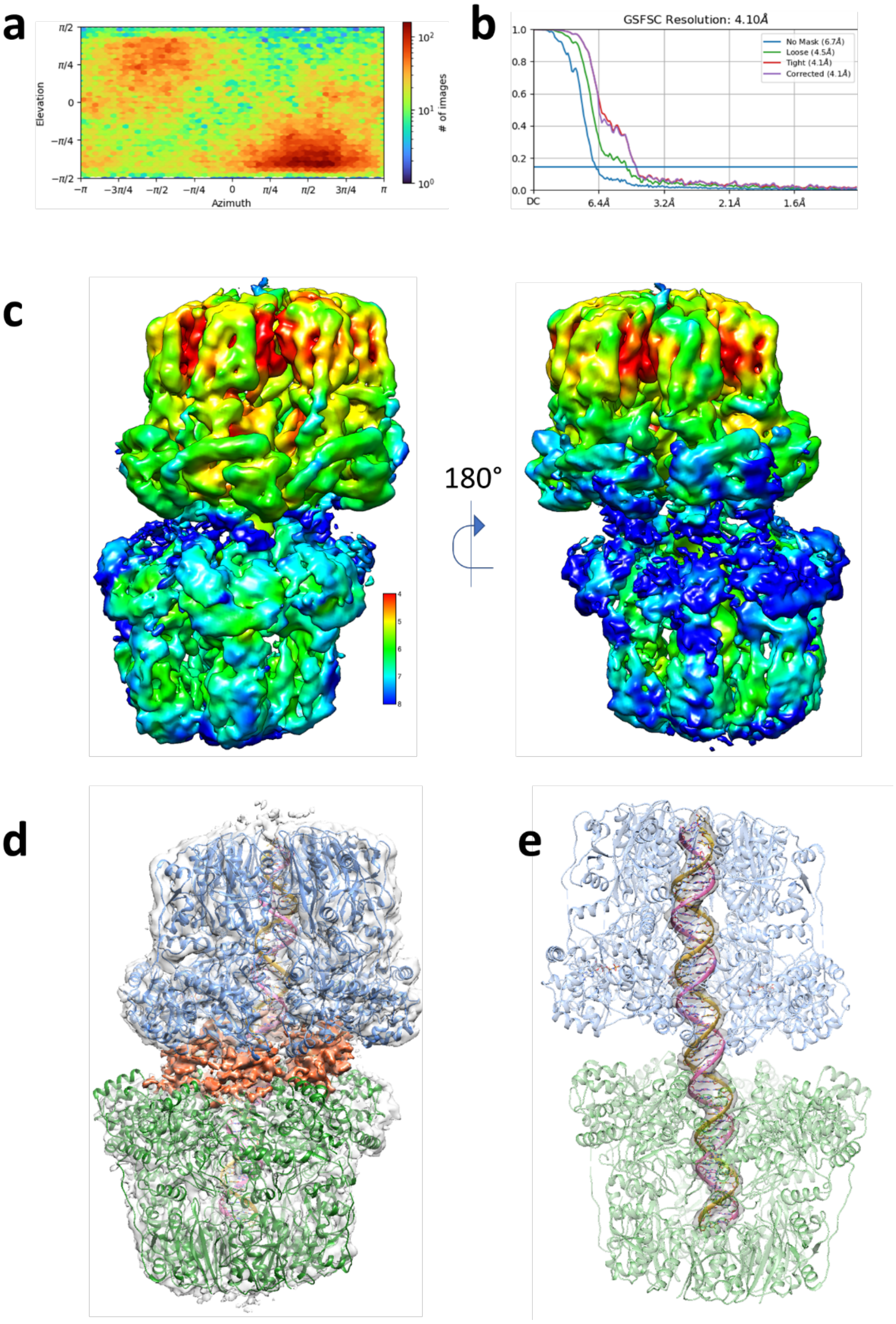

**Supplementary Figure 7: CryoEM data collection, processing and modeling of ComM dodecamer.** (**a**) Particle orientation distribution of ComM dodecamer. (**b**) Fourrier Shell Correlation (FSC) map obtained from half-maps in Cryosparc. (**c**) Local-filtered map of ComM dodecamer, at level 0.04, coloured by local resolution(in Å). (**d**) CryoEM density of ComM dodecamers, shown with 70% transparency at level 0.04. Two ComM hexamers are fitted in the density, shown in blue and green ribbons, alongside a dsDNA molecule in magenta and golden encompassing the two hexamers. In between the fitted hexamers, an additional density is observed in the position equivalent to the expected ZF domain, highlighted in orange. (**e**) Density corresponding to the 47 bp dsDNA encompassing both hexamers in the ComM dodecamer. Density for the DNA is shown at 50 % transparency while the rest of the density is hidden. Fitted dsDNA is depicted as ribbons in magenta and golden, with bases represented as sticks and colored by heteroatom. ComM hexamers are shown as blue and green ribbons, with 70% transparency.

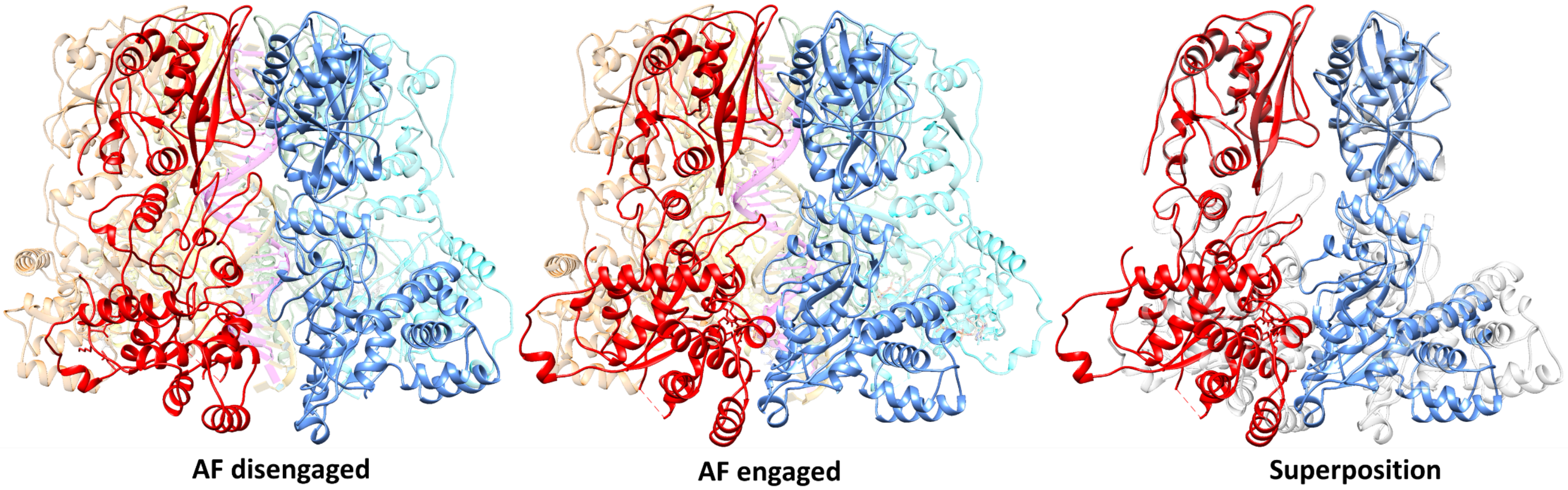

**Supplementary Figure 8: Comparison between two ComM hexamer populations.** On subpopulation 1 (left panel – EMDB 19574), A and F subunits (red and blue, respectively) are disengaged in relation to each other. In subpopulation 2 (central panel – EMDB 19575), A and F subunits are closely engaged, with consequent disengagement of F from E (cyan). The right panel shows superposition of subunits A and F, aligned on the Lon domains, for subpopulation 1 (gray ribbons) and subpopulation 2 (coloured ribbons).

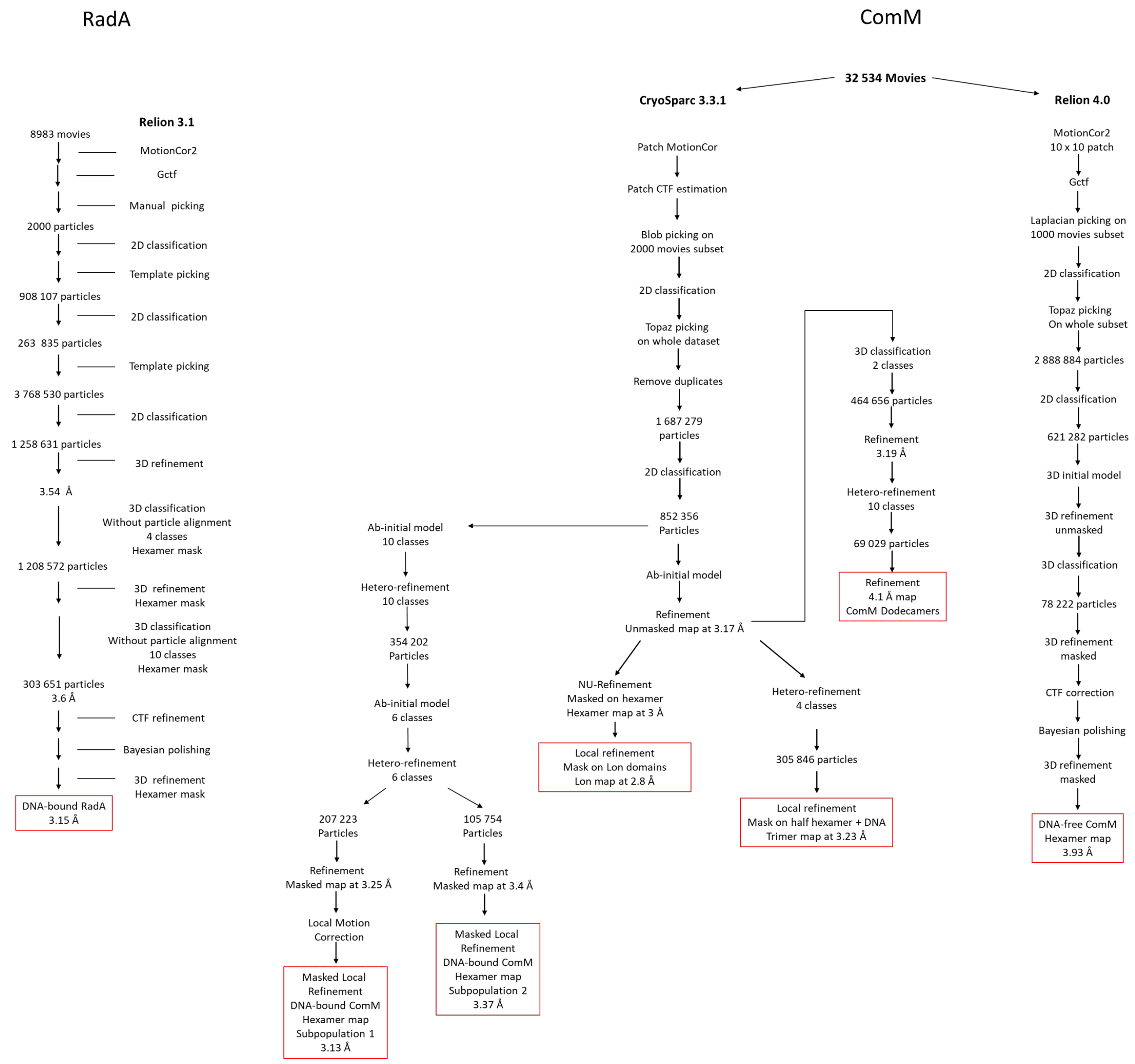

**Supplementary Figure 9: CryoEM processing workflow for RadA and ComM.**
